# mRNA-based tuberculosis vaccines BNT164a1 and BNT164b1 are immunogenic, well-tolerated and efficacious in rodent models

**DOI:** 10.1101/2025.09.30.679428

**Authors:** Neha Agrawal, Louis S. Ates, Stefan Schille, Anuhar Chaturvedi, Janina Vogt, Natasa Vukovic, Annette B. Vogel, Jan Diekmann, Mustafa Diken, Uğur Şahin

## Abstract

We designed and preclinically tested two mRNA-LNP-based vaccine candidates to protect against tuberculosis (TB). BNT164a1 and BNT164b1 encode the same eight *Mycobacterium tuberculosis* (*Mtb*) antigens expressed across different infection stages: Ag85A, Hrp1, ESAT-6, RpfD, RpfA, HbhA, M72, and VapB47. BNT164a1 utilizes nucleoside-unmodified mRNA, while BNT164b1 utilizes N1-methyl pseudouridine-modified mRNA. Prime-boost immunization with BNT164 candidates elicited antibody and/or T-cell responses against all antigens in three mouse strains (C57BL/6, BALB/c, and HLA-A2.1/DR1 humanized mice). The candidates demonstrated favorable safety profiles in a rat toxicity study and significantly reduced bacterial burdens of two *Mtb* strains in murine aerosol challenge models. BNT164 protection correlated with granuloma infiltration by CD8^+^ T cells with memory precursor phenotypes. In conclusion, BNT164a1 and BNT164b1 were immunogenic, well tolerated and efficacious in preclinical models and are the first mRNA-based TB vaccines to enter phase I/II clinical trials (NCT05537038, NCT05547464).

## INTRODUCTION

Tuberculosis (TB) is the leading cause of death by a single infectious agent, *Mycobacterium tuberculosis* (*Mtb*), causing approximately 1.25 million deaths in 2023^1^. The standard antibacterial therapy for TB is lengthy, includes multiple antibiotics with a variety of associated side effects, and can lead to emergence of drug-resistant bacteria. The only licensed TB vaccine is Bacillus Calmette-Guérin (BCG), a live attenuated vaccine derived from *M. bovis*, which is routinely administered in endemic countries. While BCG provides protection from severe forms of TB (e.g., meningeal and disseminated) in childhood, it has limited to no protective efficacy against pulmonary TB in adolescents and adults, and is contraindicated in immunocompromised individuals^2,3^. In addition to representing the highest burden of disease, pulmonary TB in adolescents and adults is responsible for the majority of TB transmission^4^. Thus, a well-tolerated, efficacious, and durable vaccine is urgently needed to prevent TB disease in these populations.

*Mtb* infection manifests as a spectrum from asymptomatic to symptomatic disease, with outcomes ranging from bacterial clearance to dissemination and death^5^. Even within one individual, highly heterogenic lung lesions co-exist independently^5,6^. This heterogeneity is dictated by an interplay between the host immune system and metabolic adaptations of the bacteria, resulting in active replication, non-replication and reactivation^6^. Preventing TB disease in adolescents and adults therefore likely requires a vaccine that targets multiple antigens expressed throughout *Mtb* infection states.

The lack of correlates of protection in humans further complicates vaccine development^7^. Preclinical data suggest antigen-specific CD4^+^ Th1 cells, and IFNγ and TNFα secretion are involved in suppressing *Mtb* infection^8,9^. Indeed, HIV infection^6^, genetic deficiencies in IFNγ pathways^10^, and anti-TNFα therapy^11^, increase patients’ risk of developing TB. However, an increasing body of evidence indicates that T follicular helper-like cells, Th1/Th17 cells, MHC restricted or non-restricted *Mtb*-specific CD8^+^ T cells and many other immune cell (sub)types also play protective roles in TB^8,9,12^. Protection conferred by humoral and B-cell responses is also increasingly studied, although the scope and mechanisms remain to be determined^13–15^.

We hypothesized that mRNA-based platforms would be appropriate for TB vaccine development, as these can induce robust cellular (CD4^+^ and CD8^+^) and antibody responses^16^. mRNA vaccines developed against COVID-19 may additionally provide valuable platform learnings that can be applied to TB vaccine development^17–19^. Furthermore, mRNA allows the encoding of multiple antigens and can be rapidly scaled up for clinical vaccine production^20^. Different mRNA chemistries have been previously investigated in humans: 1) N1-methyl pseudouridine-containing modified RNA (modRNA) which is used in two currently licensed mRNA vaccines against COVID-19 and 2) nucleoside-unmodified RNA (uRNA) which was explored in early clinical trials of COVID-19 vaccine development (NCT04380701) and currently in oncology^21,22^. These two RNA platforms differ in the way they interact with pattern recognition receptors^23^, thus, in their adjuvant activity potentially affecting the vaccine effectiveness^24–26^. Although these data obtained in other indications cannot be inferred to predict immunogenicity or efficacy of a TB vaccine, the promising platform characteristics led us to explore both platforms in a TB specific setting.

Here, we designed two multi-antigen TB vaccine candidates–BNT164a1 and BNT164b1– based on mRNA-lipid nanoparticle (mRNA-LNP) platforms. The candidates encode the same eight *Mtb* antigens and only differ in the RNA chemistry (uRNA for BNT164a1 or modRNA for BNT164b1). The aim of this study was to evaluate the immunogenicity, safety, and efficacy of BNT164a1 and BNT164b1 in preclinical models. Immunogenicity was tested in three mouse strains with different MHC backgrounds (C57BL/6, BALB/c, and HLA-A2.1/DR1 humanized mice). Safety was evaluated in a Good Laboratory Practice (GLP)-compliant toxicity study in rats, while efficacy against two different *Mtb* isolates was tested in low-dose aerosol challenge models in C57BL/6 mice.

## RESULTS

### Vaccine design

A multi-antigen strategy was adopted for the design of BNT164a1 and BNT164b1 to ensure broad immune coverage. Each candidate contains eight *Mtb* antigens (Ag85A, ESAT-6, VapB47, Hrp1, RpfA, RpfD, M72 and HbhA) distributed among four mRNAs, where each mRNA encodes a fusion of two antigens (Fig. 1A). These antigens are expressed during different stages of infection, and are immunogenic and protective against TB in different models (Table S1). Six of these antigens (Ag85A, ESAT-6, VapB47, Hrp1, RpfA, and RpfD) are part of a cytomegalovirus vector vaccine candidate that has shown protection in non-human primates (NHP) (41% sterilizing immunity)^27^, while M72 (fusion of the *Mtb* antigens PepA/Mtb32A and PPE18/Mtb39A) has shown efficacy against TB in a phase IIb clinical trial^28^. HbhA, a surface protein involved in extrapulmonary dissemination of *Mtb*^29^, was included as a potential antibody and T-cell target. Because of the high HLA diversity of populations TB endemic regions, all antigens were encoded as full-length proteins, excluding N-terminal methionines and predicted signal peptides (Table S1), to provide multiple CD4^+^ and CD8^+^ T-cell epitopes.

**Fig. 1.**
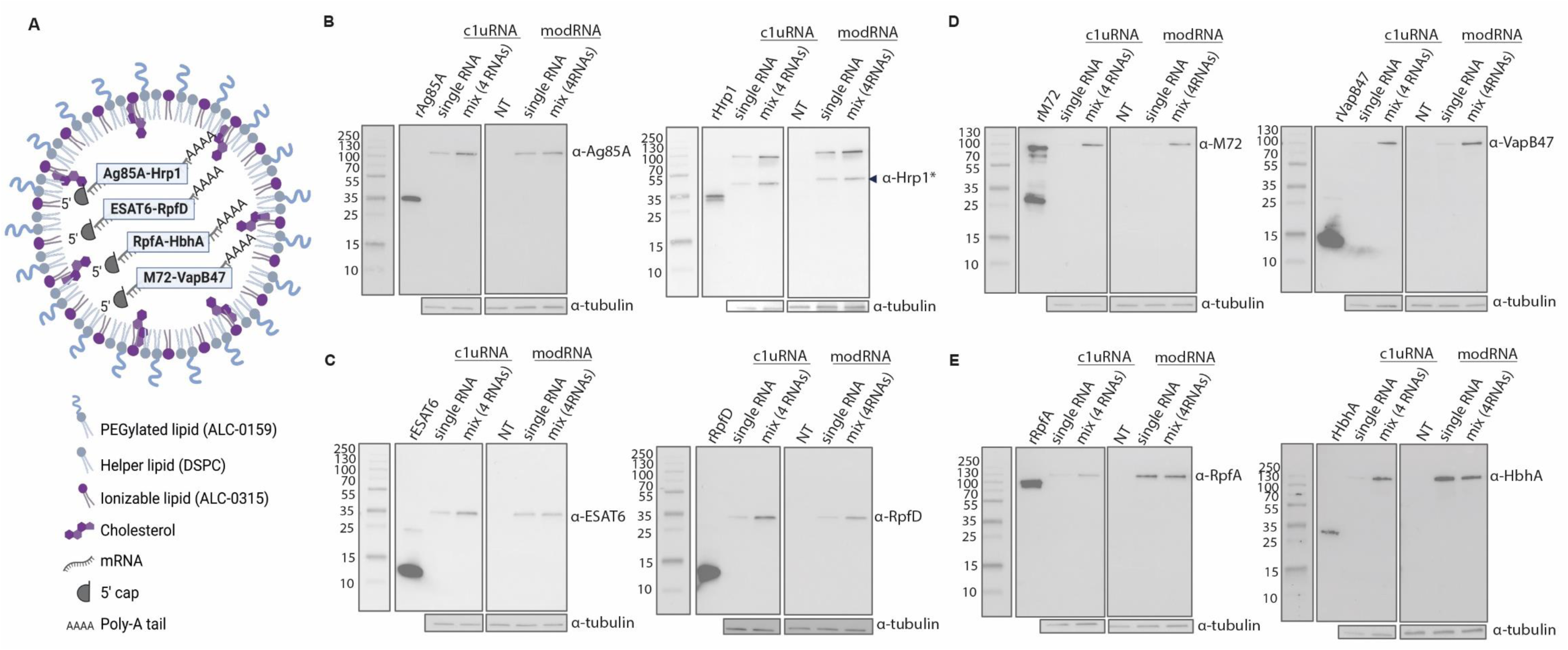
mRNAs encoding fusion *Mtb* antigens are successfully translated into corresponding proteins *in vitro.* (A) Schematic illustration of BNT164 vaccine candidates. Four mRNAs, each encoding a fusion of two *Mtb* antigens, were formulated as lipid nanoparticles. (B-E) HEK293T cells were transfected with BNT164 mRNAs either as single mRNA (0.25 µg/mL) or manually generated mixture of the four mRNAs (1 µg/mL) using RiboJuice^TM^ transfection kit. Non-transfected cells (NT) were used as a negative control, and respective recombinant (r) proteins as positive controls. Protein expression was assessed by western blot using antigen-specific antibodies. Representative western blots are shown. Expected molecular weights: (B) Ag85A–Hrp1: 50 kDa; (C) ESAT-6–RpfD: 28 kDa; (D) M72–VapB47: 86 kDa; and (E) RpfA–HbhA: 120 kDa. Black arrow indicates Ag85A-Hrp1 monomer. Loading control: αtubulin. *Anti-Hrp1 antibody detected both monomeric and dimeric Ag85A-Hrp1 fusion protein.

### mRNA-encoded antigens are expressed *in vitro*

To confirm successful protein expression of BNT164a1 and BNT164b1 mRNA constructs in human cells, we transfected HEK293T cells with either individual mRNAs (each containing two antigens) or with a mix of all four mRNAs per vaccine candidate and performed western blot analysis. All eight mRNA-encoded *Mtb* antigens were detected in HEK293T cell lysates at expected molecular weights (calculated for fusion proteins) (Fig. 1B-E). Anti-Hrp1 detection revealed the presence of two bands with molecular weights corresponding to monomeric and dimeric Ag85A-Hrp1 fusion protein. Dimerization of Hrp1 has been previously reported^30^, and our data indicates that it was not affected by fusion to Ag85A. The monoclonal Ag85A antibody detected only the dimeric version of Ag85A-Hrp1. Protein expression did not decrease for any of the constructs when mRNA was delivered in a mix compared with individual mRNAs, suggesting no major translational interference *in vitro*. Taken together, the mRNAs used in BNT164 candidates were successfully translated into corresponding *Mtb* fusion proteins *in vitro*.

### BNT164 candidates elicit cellular and humoral immunogenicity in mice

We assessed BNT164 immunogenicity by analyzing T- and B-cell responses in a series of experiments in different mouse strains, following the immunization scheme shown in Fig. 2A.

**Fig. 2.**
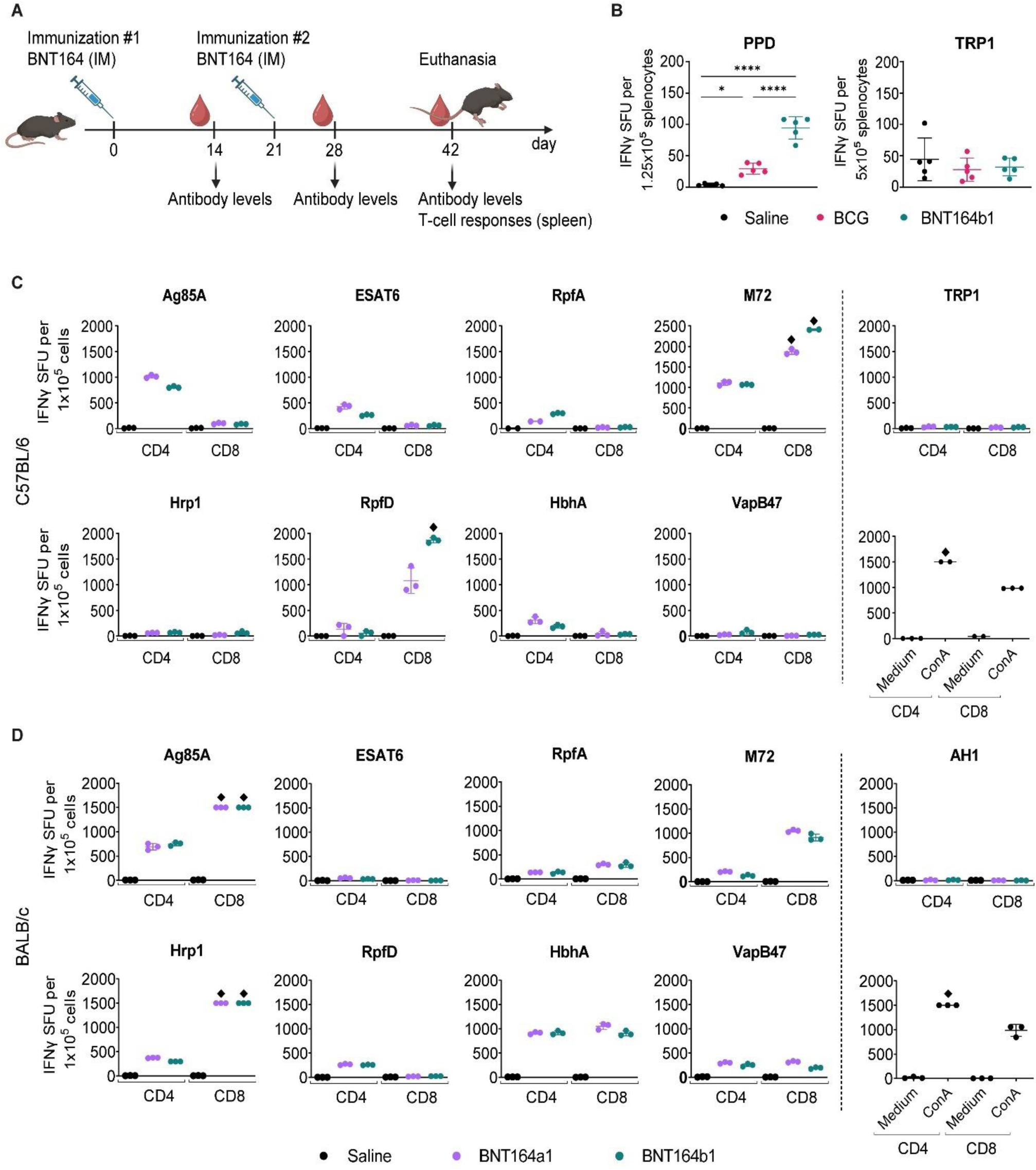
BNT164 vaccine candidates induced T-cell responses against PPD and all target *Mtb* antigens in mice. (A) Design of immunogenicity studies performed with BNT164 candidates, unless indicated otherwise. C57BL/6 mice (B-C) or BALb/c mice (D) were immunized on Day 0 with a saline control (IM) or 4 µg BNT164a1/b1 (IM) on Days 0 and 21, or with ∼10^6^ colony-forming units BCG (SC; only in (B)). (B) Splenocytes were isolated on Day 42 and stimulated with purified protein derivative (PPD) or TRP1 (non-specific peptide) and responses were assessed by IFNγ ELISpot assay after ∼18 hours incubation. Group mean values are indicated by horizontal bars (± standard deviations), means from individual mice (n=5/group, measured in duplicates) are depicted as circles. One-way ANOVA with Tukey’s multiple comparisons test was performed: *p < 0.05; ****p < 0.0001. (C–D) Splenocytes were isolated on Day 42. CD4^+^ and CD8^+^ T cells were magnetically sorted from splenocyte pools from each group (n=5/group) and stimulated with individual *Mtb* antigens or non-specific peptides (TRP1, AH1 respectively). Media only and concanavalin A (con A) were assay controls. Bone marrow derived dendritic cells from non-vaccinated mice (unrelated cohort of naïve C57BL/6 or BALB/c mice) were co-cultured with T cells as antigen-presenting cells. Responses were assessed by IFNγ ELISpot assay after ∼18 hours incubation. Circles represent duplicate/triplicate measurements; horizontal bars represent group means ± standard deviation. The rhombus symbol indicates the conditions where the upper limit in number of spots that can be correctly counted was reached (∼1500 spots). Statistical analysis was not performed to evaluate the differences, since the data are technical replicates of pooled samples within each treatment group. IM = intramuscular; SC = subcutaneous; SFU = spot-forming unit.

#### T-cell responses

For a preliminary assessment of T-cell immunogenicity, we compared Day 42 splenic T-cell responses induced by BCG (immunization: subcutaneous [SC], Day 0) and BNT164b1 (immunization: intramuscular [IM], Days 0 and 21) in C57BL/6 mice, upon *ex vivo* restimulation with purified protein derivative (PPD) in an ELISpot assay. While both BCG and BNT164b1 induced a significantly higher number of IFNγ-secreting T cells compared with saline, BNT164b1 was superior to BCG (Fig. 2B). Stimulation with unrelated peptide TRP1 served as a control for non-specific responses and induced a low number of IFNγ-secreting cells in all test groups (Fig. 2B).

We next analyzed individual T-cell responses against each antigen encoded by BNT164 candidates in mice immunized with BNT164a1, BNT164b1, or saline. IFNγ secretion from total splenocytes or purified CD4^+^ or CD8^+^ T cells was measured using ELISpot assay three weeks after the boost (Day 42). BNT164a1 and BNT164b1 raised CD4^+^ or CD8^+^ T-cell responses, or both, against each of the eight *Mtb* antigens in at least one of the mouse strains (C57BL/6 or BALB/c) (Fig. 2C-D). T-cell responses against Ag85A, Hrp1 and HbhA were generally higher in BALB/c mice than in C57BL/6, whereas responses against M72 and RpfD were higher in C57BL/6 mice. Stimulation with non-specific peptides TRP1 and AH1, or medium only, induced minimal CD4^+^ or CD8^+^ T-cell responses (Fig. 2C-D). To understand potential immunogenicity for certain human HLA alleles, we next used HLA-A2.1/DR1 humanized mice in which endogenous MHC molecules are knocked out and human HLA-A2.1 and HLA-DR1 are expressed instead^31^. In this model, BNT164 candidates also raised antigen-specific T-cell responses against all antigens (Fig. S1). ELISpot analysis using total splenocytes reflected the results obtained with purified CD4^+^ and CD8^+^ splenic T cells in all three mouse models (Fig. S2).

For a more detailed analysis of T-cell responses, we quantified the percentages of CD4^+^ and CD8^+^ cells producing IFNγ, IL-2, or TNFα, or all three cytokines, as well as T cell memory subsets, via flow cytometry three weeks after the boost (Day 42). Following splenocyte stimulation with a pool of peptides against all 8 *Mtb* antigens, we observed significantly higher percentages of single cytokine-secreting CD4^+^ and CD8^+^ T cells and polyfunctional T cells (IFNγ^+^/IL-2^+^/TNFα^+^) in C57BL/6 mice immunized with BNT164 candidates compared with saline (Fig. S3). While there was no difference between candidates in elicited CD4^+^ responses, BNT164a1 induced a significantly higher percentage of CD8^+^ cytokine-secreting T cells than BNT164b1 (Fig. S3). Analysis of T cell memory subsets revealed that BNT164 candidates induced both memory precursor effector cells (MPEC: cytokine^+^/CD127/^+^KLRG1^-^/CD62L^-^) and long-lived memory precursor (LLMP: cytokine^+^/CD127^+^/KLRG1^-^/CD62L^+^) cells (Fig. S3). Similar to observations with cytokine-producing T cells, BNT164a1 induced significantly higher percentages of CD8^+^ memory precursor cells compared with BNT164b1.

Taken together, BNT164 candidates induced cellular immunity against the selected *Mtb* antigens in multiple tested mouse strains following prime-boost dosing.

#### IgG responses

We assessed BNT164-induced humoral responses in sera of C57BL/6, BALB/c and HLA-A2.1/DR1 humanized mice using ELISA. In C57BL/6 (Fig. 3A) and BALB/c (Fig. S4A) mice, prime-boost immunization with BNT164a1 or BNT164b1 elicited IgG responses against all *Mtb* target antigens, except ESAT6 and RpfD. IgG responses were mostly detectable by Day 14 post-prime dose and titers increased until the end of experiment on Day 42. We detected comparable levels of IgG antibodies against Ag85A, RpfA, and Hrp1 in both mouse models, but observed some strain-specific differences in IgG responses against M72 (higher in C57BL/6) and HbhA (higher in BALB/c). As expected, in HLA-A2.1/DR1 humanized mice, BNT164a1 and BNT164b1 immunization induced only marginal IgG responses against a few antigens (Fig. S4B).

**Fig. 3.**
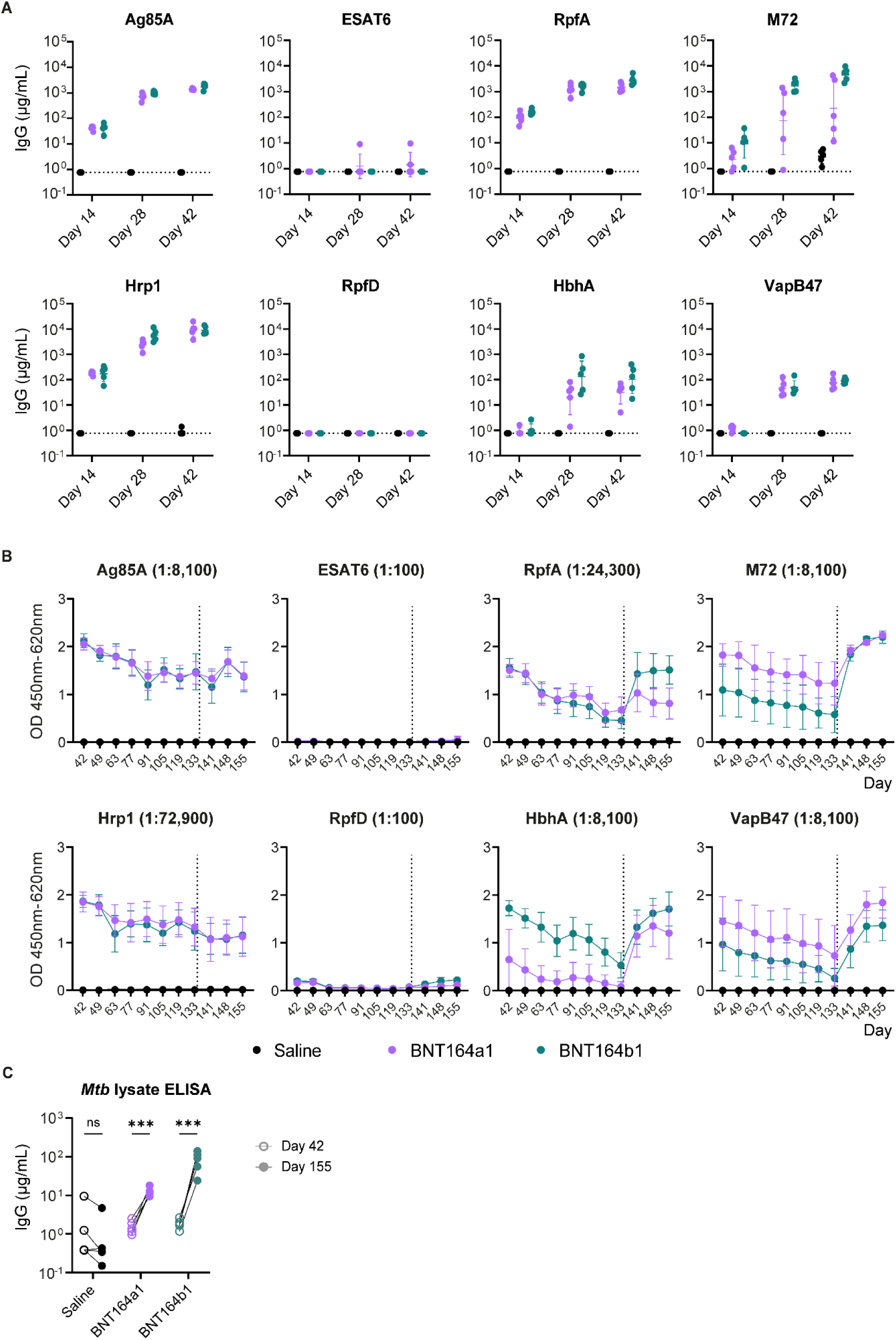
BNT164 vaccine candidates induced IgG responses against majority of vaccine antigens in C57BL/6 mice. C57BL/6 mice received IM injection of 4 µg BNT164a1, 4 µg BNT164b1, or a saline control on (A) Days 0 and 21, or (B– C) Days 0, 21 and 134. Sera isolated on indicated days was used to measure (A–B) antigen-specific IgG antibodies by ELISA, using recombinant *Mtb* proteins, or (C) anti-*Mtb* IgG antibodies by ELISA, using *Mtb* H37Rv lysate. (A–B) Group mean values are indicated by horizontal bars (± standard deviations); means from individual mice (n=5/group, measured in duplicates) are depicted as circles. Horizontal dashed line in (A) indicates lower limit of detection (LLOD). Vertical dashed line in (B) indicates the timing of third immunization. (C) Means from individual mice (n=5/group, measured in duplicates) are depicted as circles; lines connect the same mouse on different days. Differences between days for each group were compared with ratio-paired t tests corrected for multiple comparisons by false discovery rate calculation using Benjamini, Krieger, Yekutieli methodology: ns = q > 0.05; *** = both q and p < 0.001. IM = intramuscular; *Mtb* = *Mycobacterium tuberculosis*.

Next, we investigated durability and boostability of IgG responses in C57BL/6 mice immunized with two BNT164a1 or BNT164b1 doses (as above). Biweekly monitoring revealed a modest gradual decrease in antibody titers from Day 42 to Day 133 (Fig. 3B). For antigens where the titers decreased the most (RpfA, HbhA, M72 and VapB47), the third BNT164a1 or BNT164b1 immunization on Day 134 successfully boosted the responses to similar or higher levels than observed on Day 42 (Fig. 3B).

Finally, to confirm that BNT164-induced antibodies can bind to *Mtb* antigens in their native conformation, we performed an *Mtb* H37Rv lysate ELISA with sera of immunized mice. Of note, *Mtb* lysate likely does not include Hrp1, VapB47, RpfA and RpfD, since these antigens are not expressed in general culture conditions. IgG binding to *Mtb* lysate was observed at relatively low levels at Day 42 for all BNT164-immunized animals and one saline-injected animal (Fig. 3C). The IgG response against the *Mtb* lysate increased significantly between Days 42 and 155 in both BNT164a1 (8-fold) and BNT164b1 (40-fold) groups. These data suggest that the third dose of BNT164 increased the affinity of induced antibodies, since binding to *Mtb* lysate with limited antigen availability was markedly increased, while binding to recombinant proteins remained comparable between Days 42 and 155.

### BNT164 candidates show a favorable safety profile in rats

To evaluate the safety profiles of BNT164a1 and BNT164b1, we performed a GLP-compliant toxicity study in which rats received four weekly IM injections of BNT164a1 (10 µg), BNT164b1 (10 µg or 30 µg), saline, or BNT164 non-translatable control (30 µg, modRNA). Toxicity was assessed at two time points: the main study group was sacrificed two days after the last immunization, while the recovery group was sacrificed 3 weeks after the last immunization to study the reversibility of vaccine-induced changes (Fig. 4A).

**Fig. 4.**
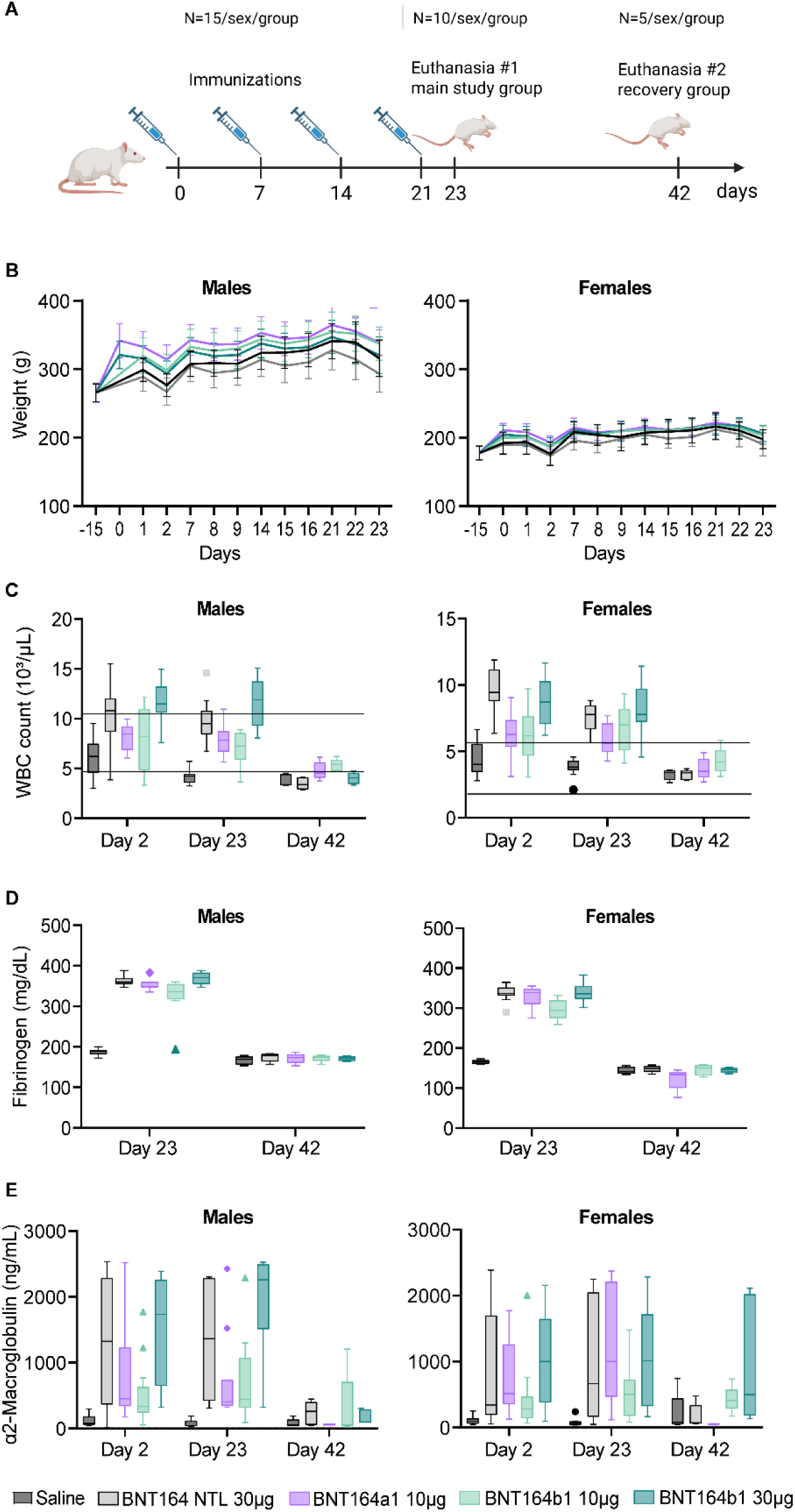
BNT164 candidates were well tolerated in a repeat-dose toxicity study in Wistar Han rats. Wistar Han rats were injected IM with saline, BNT164 non-translatable control (NTL, 30µg), BNT164a1 (10µg), or BNT164b1 (10µg or 30µg), on Days 0, 7, 14, and 21. Main study group was sacrificed on Day 23 (n=10), while the recovery group was sacrificed on Day 42 (n=5). Some samples were unsuitable for hematological analysis following coagulation and/or following unreliable instrumental data and are indicated in the method section. (A) Schematic representation of the study. (B) Animal weight (mean ± standard error of the mean). (C–E) Laboratory findings plotted with Tukey box plots showing group medians (middle line), 25th and 75th percentiles (box), upper and lower adjacent values (whiskers), and outliers (symbols). Horizontal lines in (C) represent the range of historical values of white blood cells (WBC) for this strain at study site. IM = intramuscular.

We did not observe any vaccine-related systemic clinical signs or mortalities, nor vaccine-related changes in the body weight (Fig. 4B) or urinalysis. In males, there were no remarkable changes in body temperature throughout the study, while an increase in body temperature was observed in females 24 hours after the first injection of BNT164a1, and 24 hours after the third injection of 30 µg BNT164b1 (Fig. S5A). Macroscopic observations of the injection sites did not reveal signs of redness, swelling, scabbing or other macroscopic alterations. However, histological examination revealed muscle fiber damage and inflammatory reactions related to BNT164 administration (data not shown) that partially or completely resolved by Day 42. On Days 2 and 23, the BNT164 candidates induced increased levels of white blood cells, neutrophils, large unstained cells, and acute phase proteins (fibrinogen, α2-Macroglobulin and α-1-acid glycoprotein), indicating expected inflammatory responses to a vaccine (Fig. 4C-E, Fig. S5B-D). The changes were transient: all values returned to normal by Day 42, except for α-2-Macroglobulin levels in some animals treated with BNT164b1 (both doses), and α-1-acid glycoprotein levels in some males treated with BNT164a1 and some females treated with 10 µg BNT164b1.

All observed changes were dose-dependent, as the animals vaccinated with 30 μg BNT164b1 showed a higher increase of inflammatory parameters compared with animals vaccinated with 10 μg BNT164b1. Animals injected with non-translatable control (NTL) also showed an increase in inflammatory parameters (comparable to BNT164b1 at the same dose), indicating that the effects were formulation-dependent rather than antigen-specific. Overall, BNT164 candidates were well tolerated and induced expected inflammatory reactions that mostly resolved within 3 weeks of the last dose.

### BNT164 candidates reduce the bacterial burden in *Mtb*-challenged mice

To assess the efficacy of BNT164 candidates, we used a low dose aerosol *Mtb* challenge model with two different *Mtb* strains: The reference strain H37Rv (genetic lineage L4) and hypervirulent HN878 (genetic lineage L2). Both L2 and L4 are globally dispersed^32^ and while genetically very similar (>90% sequence identity), show phenotypic differences in antigenicity, pathogenesis and vaccine protection^33,34^. BNT164-immunized or saline-injected C57BL/6 mice were exposed to ∼100 colony-forming units (CFUs) of H37Rv or HN878 *Mtb* at week eight of the experiment, and the bacterial load in lung (site of infection) and spleen (extrapulmonary site) was determined 30 days post-infection (dpi) for both strains, and 60 dpi only for H37RV (Fig. 5A). Non-translatable BNT164 modRNA formulated in LNP (NTL) and saline were used as controls.

**Fig. 5.**
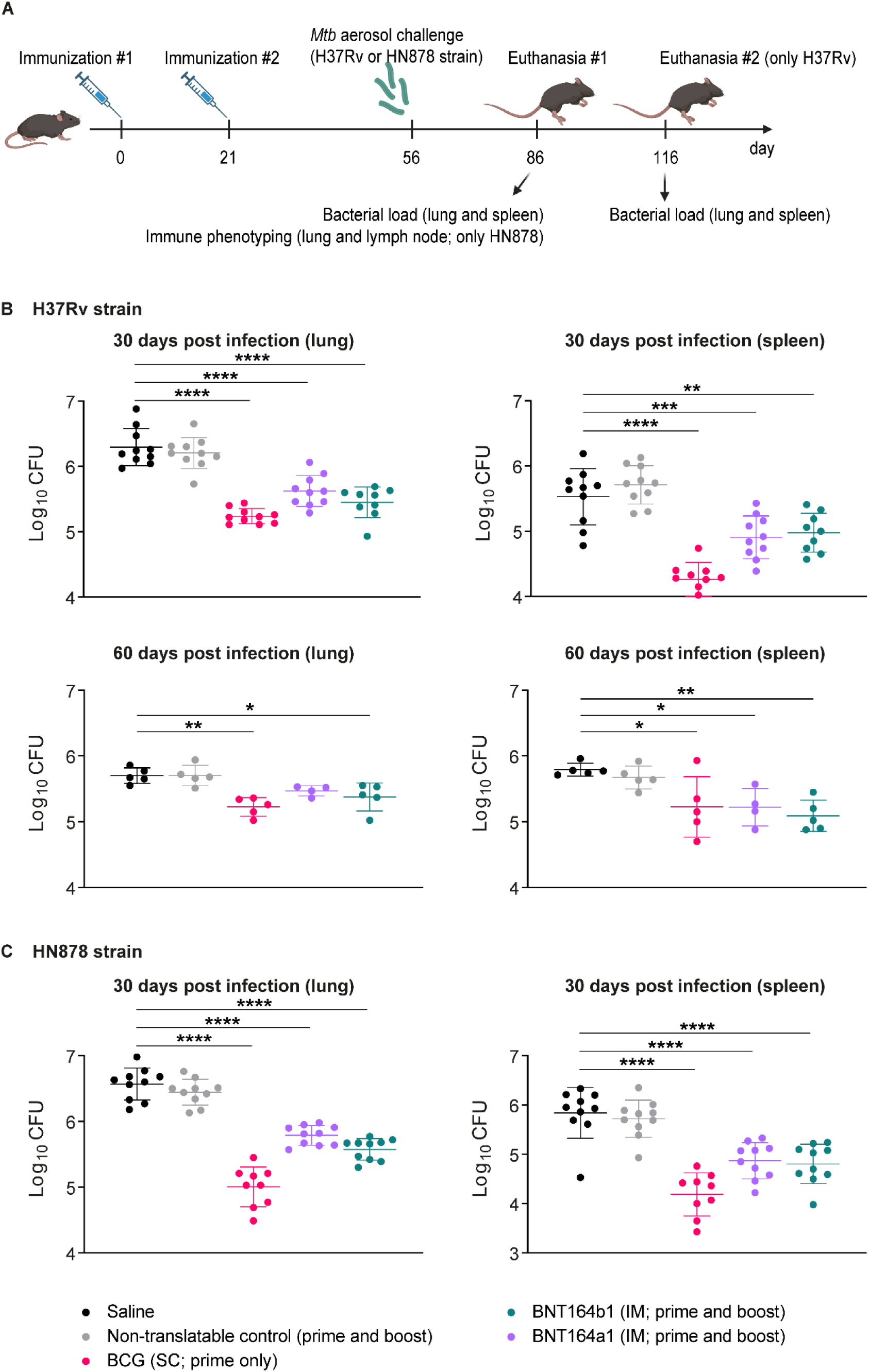
BNT164 candidates significantly reduced the bacterial load in *Mtb*-challenged mice. C57BL/6 mice were injected with saline (IM, days 0 and 21), BCG (SC 10^6^, Day 1), or mRNA-LNP (IM 4 µg, Days 0 and 21). On Day 56, mice were aerosol-challenged with ∼100 CFUs of *Mtb* H37Rv or HN878. The left lobe of the lung and the spleen were harvested, processed into a homogenate, serially diluted onto 7H11 agar, and cultured to enumerate viable bacteria on Day 86, 30 days post-infection (B top and C; n = 9–10) or Day 116, 60 days post-infection (B bottom; n = 4–5). (A) Schematic representation of the study. (B–C) CFUs in lungs (left) and spleen (right). Group mean values are indicated by horizontal bars (± standard deviations); individual mouse values are depicted as circles. One-way ANOVA with Dunnett’s multiple comparisons test between the saline-treated group and all other groups was performed: * = p < 0.05; ** = p < 0.01, *** = p < 0.001, **** = p < 0.0001. CFU = colony forming unit; IM = intramuscular; LNP = lipid nanoparticle; *Mtb = Mycobacterium tuberculosis*; SC = subcutaneous.

In H37Rv-infected mice, BNT164a1 and BNT164b1 significantly reduced bacterial burden compared with saline (∼0.55–0.85 log reduction) in both lung and spleen 30 dpi (Fig. 5B). At 60 dpi, lung CFUs remained comparable to 30 dpi values in all test groups except saline and NTL control groups, where CFU values declined (Fig. 5B). Also, all test groups showed a slight increase in splenic bacterial load at 60 dpi compared with 30 dpi (Fig. 5B). Mice immunized with NTL control did not have significant reductions in bacterial burden compared to saline group at either time point, indicating that BNT164 candidates reduced the bacterial burden predominantly via antigen-specific immunity (Fig. 5B). Mice immunized with BCG served as a positive control group and showed ∼1 log reduction of viable bacteria compared with saline 30 dpi (Fig. 5B). In HN878-infected mice, BNT164 candidates also significantly reduced the bacterial burden in lung and spleen 30 dpi compared with saline (∼0.8–1 log reduction) (Fig. 5C).

To assess the effect of BNT164 vaccination on pathology of *Mtb* HN878 infection, lung sections were evaluated histologically (Fig. S6). While all groups had structured granulomas, they differed in the presence of acid-fast bacteria. In the saline- or NTL-injected groups, >80% of granulomas contained acid-fast bacteria, whereas this was significantly lower (∼50%) for BCG and BNT164-immunized groups (Fig. 6A), in line with the bacterial burden obtained by CFU counting (Fig. 5). This reduction in bacterial burden in the groups immunized with BCG or BNT164 candidates negatively correlated with lymphocyte infiltration (r=-0.7785; p<0.0001) (Fig. 6B). In contrast, lungs from groups injected with saline or NTL showed high presence of macrophages and neutrophils. While macrophage presence was similar across all groups (data not shown), no neutrophils could be detected in the sections from animals immunized with BNT164 candidates (Fig. 6C).

**Fig. 6.**
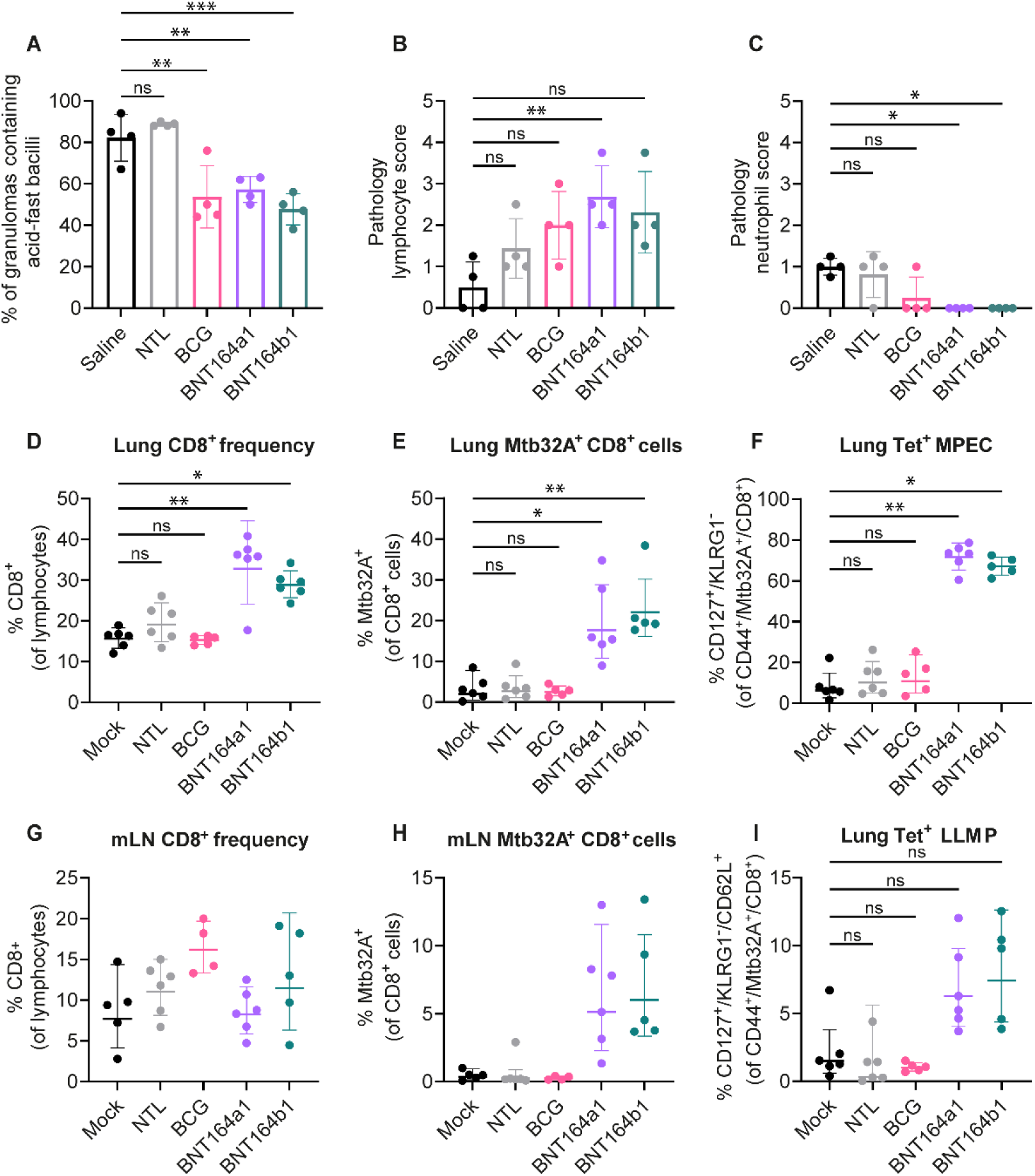
BNT164-induced protection correlates with antigen-specific CD8+ T cells with a memory precursor phenotype in the lungs. Right lung lobes of the same mice depicted in figure 5, infected with *Mtb* HN878, were harvested at the time of sacrifice 30 days post infection. For four mice per group these lungs were paraffin embedded, stained with acid-fast (A) or H&E (B–C) staining. A pathologist blinded for group identities scored the percentage of granulomas containing acid-fast bacteria (A) and scored the immune cell composition within granulomas by multiplying the granuloma severity score by cell composition (further detailed in methods) (B–C). Right lung lobes (D-F, I) and mLNs (G–H) of the remaining 5–6 mice in each group were homogenized and analyzed by flow cytometry. All plots were analyzed by ANOVA, and if significant (p < 0.05), tested for multiple comparisons of all groups against the saline control. (A) ordinary one-way ANOVA, followed by Dunnet’s test. (B–I) Kruskal-Wallis test followed by Dunńs test, except for G–H where Kruskal-Wallis p > 0.05. ns = p > 0.05; *p < 0.05; **p < 0.01, ***p < 0.001; LLMP = Long-lived memory precursor; mLN = mesenteric lymph node; MPEC = Memory precursor effector cell.

In summary, BNT164 vaccine candidates significantly reduced *Mtb* burden following aerosol challenge in mice with two different *Mtb* lineages. Protection induced by BNT164 candidates was associated with lymphocytic infiltration into the lungs.

### BNT164 candidates induce CD8^+^ T cells with memory precursor phenotypes

In parallel to the pathological assessment, small lung lobes from the remaining mice were analyzed by flow cytometry using a lymphocyte phenotyping panel (Fig. S7A). Although tetramers in the C57BL/6 background are very limited, the panel included one class I tetramer derived from Mtb32A (a subcomponent of M72) and one ESAT-6 specific class II tetramer. Consistent with microscopical findings, the BNT164-immunized groups showed a higher (although not statistically significant) total number of lung lymphocytes than negative control groups (saline- or NTL-injected) (Fig. S7B). This was not driven by increased frequencies of total or antigen-specific CD4 cells (Fig. S7C and F), but rather a significant increase (∼2-fold) in CD8^+^ T cell frequencies (Fig. 6D). A large proportion (∼20%) of these CD8^+^ cells were activated (CD44^+^) and antigen-specific as shown by the Mtb32A tetramer (Fig. 6E).

We further characterized the CD44^+^/Mtb32A tetramer^+^/CD8^+^ T cells to distinguish between short-lived effector cells (SLEC; CD127^-^/KLRG1^+^) and MPEC cells (CD127^+^/KLRG1^-^), further defined as LLMP cells (CD127^+^/KLRG1^-^/ CD62L^+^). Approximately 70% of Mtb32A tetramer^+^/CD8^+^ T cells in the lungs of BNT164-immunized groups had an MPEC phenotype, which was significantly higher than in BCG-immunized or negative control groups (∼10%) (Fig. 6F). LLMP cells were also increased in BNT164-immunized groups (Fig. 6I). Although the difference in frequencies of these cells (within CD44^+^/Mtb32A tetramer^+^/CD8^+^ T population) was not statistically significant due to low cell numbers in control groups (Fig. 6I), mean counts of Mtb32A tetramer^+^ LLMPs ranged between 8–10 for saline, BCG and NTL groups, compared to ∼791 for BNT164a1 and ∼760 for BNT64b1 (data not shown). This increase of MPEC and LLMP memory precursor cells coincided with a significant decrease in SLEC frequencies (Fig. S77D), although absolute numbers of these cells remained constant (not depicted). A trend of MPEC cell enrichment in lungs was also observed among CD44^+^/Mtb32A tetramer^-^/CD8^+^ T cells in BCG- and BNT164-immunized groups (Fig. S77E).

In contrast to the lungs, mesenteric lymph nodes (mLNs) of BNT164-immunized animals did not show an increased frequency of CD8^+^ T cells (Fig. 6G). A trend of enrichment of Mtb32a tetramer^+^ cells could be observed but was overall less pronounced than in lung tissues (Fig. 6H). In the mLNs, the MPEC population was more abundant than in lungs for all groups, but still enriched in BNT164-immunized animals (Fig. S77I). Very low number of SLECs were detected in mLNs and no enrichment of LLMP was observed (Fig. S77G-H).

Taken together, the comparative assessment of mLNs and lungs suggests that the antigen-specific CD8^+^ T cells induced by BNT164 candidates homed to the primary site of infection (lungs), exhibiting a memory precursor phenotype with hallmarks of long-lived immunity.

## DISCUSSION

Here, we describe two mRNA-LNP TB vaccine candidates, BNT164a1 and BNT164b1. Both candidates encode the same eight full-length *Mtb* antigens as fusion proteins on four mRNAs, that are successfully expressed in human cells. Prime-boost BNT164 immunization induced T-cell and/or IgG responses against each of the eight encoded *Mtb* antigens, with expected differences in type and intensity of responses between tested mouse strains. In low dose aerosol *Mtb* challenge studies in C57BL/6 mice with two different *Mtb* strains, mice vaccinated with BNT164a1 or BNT164b1 had significantly lower bacterial load in lungs and spleens than those injected with saline.

The only difference between BNT164 candidates is the mRNA platform used. Evidence suggests different adjuvant activity between these platforms: while nucleoside-unmodified RNA is considered to signal through Toll-like receptors (TLR3, TLR7 and TLR8) and exhibit strong adjuvant activity, modRNA activates other innate immunity pathways (e.g., melanoma differentiation-associated protein 5 (MDA5) signaling) and has lower adjuvant activity^23,25^. We detected higher levels of polyfunctional (IFNγ^+^/TNFα^+^/IL-2^+^) and memory CD8^+^ cells in the BNT164a1 (uRNA) group compared with the BNT164b1 (modRNA) group. This is in alignment with previous findings in a B16-OVA murine melanoma model where nucleoside-unmodified RNA induced higher frequency of polyfunctional (granzyme B^+^/IFNγ^+^/TNFα^+^) antigen-specific CD8^+^ T cells compared with modRNA, resulting in better tumor growth control^26^. However, we did not find consistent significant differences between modRNA- and uRNA-induced T-cell responses via IFNγ ELISpot, nor in T-cell phenotypes and protection following *Mtb* challenge. Platform differences between BNT164a1 and BNT164b1 may be underestimated in murine models due to a generally lower sensitivity to innate immune signals compared with humans^35^. Thus, whether the use of different mRNA platforms affects the efficacy of BNT164 candidates remains to be explored in humans.

The lack of established animal models for efficacy testing of TB vaccine candidates poses a hurdle in TB vaccine development. Although many different TB preclinical challenge models can be explored, such as different mouse backgrounds, other rodent or NHP models, different dose, route, duration and timing of the challenge, or vaccination after BCG prime or *Mtb* exposure, none of these are proven to predict human protection^7,36,37^. Therefore, we first selected the most widely used model in the field, a low dose aerosol *Mtb* H37Rv challenge in C57BL/6 mice, and showed that BNT164a1 and BNT164b1 significantly reduced the bacterial load in lungs and spleen of infected mice when compared with saline control. In the second challenge study, we extended these findings and demonstrated protection against the hypervirulent L2 isolate HN878. In C57BL/6 mice, HN878 has been reported to induce TB immune responses that more closely resemble those in humans, such as formation of granulomas^38^, granuloma-associated lymphoid tissues^39^, and a protective role for IL-17 signaling^34^. Still, our challenge studies have limitations that should be considered when interpreting the data. These include the lack of balanced host-pathogen interactions in mice, the lack of expression of hypoxia- (Hrp1 and VapB47) and resuscitation-(RpfA and RpfD) associated antigens in the investigated timeframes, and the limited MHC repertoire of C57BL/6 mice. Further BNT164 immunogenicity and efficacy testing, in naïve and Mycobacterial antigen-exposed individuals, therefore, remains to be determined in clinical trials.

Despite the limitations of the animal models utilized here, they provide encouraging findings of diverse and broad immune responses induced by BNT164 candidates. An important vaccine candidate aimed at preventing TB in adolescents and adults is the M72/AS01E subunit vaccine candidate, which showed efficacy in preventing TB disease in *Mtb*-exposed individuals (positive in the IFNγ release assay)^28,40^ and is currently being evaluated in a phase III trial (NCT06062238). M72/AS01E induced polyfunctional CD4^+^ T cells and high IgG responses, with CD8^+^ T-cell responses detected in a phase 2 immunogenicity study^41^, but not in the phase 2b efficacy trial possibly due to different timepoints investigated^28^. While the mechanisms of protection of M72/AS01E remain to be elucidated, it is encouraging that, similar to M72/AS01E, BNT164 candidates induced strong IgG and CD4^+^ responses to M72 in mice, including polyfunctional CD4^+^ cells secreting (IFNγ, TNFα and IL2)^42^. Nonetheless, expected efficacy of BNT164 candidates cannot be inferred from immunological comparisons with other vaccine candidates due to the lack of correlates of protection and different modes of action. In addition to anti-M72 responses, BNT164 candidates also elicited T-cell and antibody responses to the other *Mtb* antigens included in the vaccine. Beyond M72, we have identified durable IgG antibody responses to Ag85A, Hrp1, RpfA, HbhA, and VapB47. For the antigens where the IgG responses reduced over time, a third vaccine dose successfully boosted the titers. Furthermore, a third dose increased binding of serum IgG to *Mtb* H37Rv lysate, possibly by increasing affinity maturation. Whether these antibodies contribute to TB protection remains to be established.

In addition to CD4^+^ and IgG responses, BNT164 elicited cytokine-producing CD8^+^ T cells, for which our data suggest a potentially beneficial role in protection. In the *Mtb* challenge study, higher numbers of lymphocytes infiltrating the granulomas correlated with lower bacterial burdens. Observed lymphocyte infiltration was driven by antigen-specific CD8^+^ T cells specifically in the lungs as the primary site of infection. Furthermore, both in the immunogenicity and the challenge setting, the BNT164-induced CD8^+^ T cells exhibited memory precursor phenotypes. The contribution of CD8^+^ T-cells in TB protection is not fully elucidated, but recent data suggest important roles in the NHP model^43^ and that the interplay between CD4^+^ and CD8^+^ T cells is essential for optimal immunity in mice^44^.

BNT164a1 and BNT164b1 were well tolerated in a GLP-compliant repeat-dose toxicity study in rats, with no vaccine-related systemic clinical signs or mortalities. The candidates did induce transient dose-dependent inflammatory responses attributed to the vaccine-induced activation of immune system: increased acute phase reactants, increased counts of white blood cells, transiently increased body temperature, and microscopic inflammation at the injection site. The control group immunized with non-translatable modRNA-LNP had similar findings, indicating formulation-dependent activation of innate immunity, rather than antigen-dependent inflammation. These results are consistent with SARS-CoV-2 mRNA-LNP vaccine studies, where a similar inflammatory response was seen^45^.

In conclusion, preclinical data showed that TB vaccine candidates BNT164a1 and BNT164b1 are immunogenic, well tolerated and efficacious in rodent models, supporting their further clinical development. BNT164a1 and BNT164b1 are the first mRNA-based vaccines against *Mtb* to enter phase I/II clinical testing, where their safety and immunogenicity are being evaluated (NCT05537038, NCT05547464).

## METHODS

### Study design

The objective of this study was to assess immunogenicity, safety and efficacy of two mRNA vaccine candidates against TB in preclinical models. Statistical methods were not used to predetermine group sizes since effect sizes were unknown before testing. Only animals with an unobjectionable health status were selected for testing procedures and humane endpoints were predefined. Animals were randomized only for the toxicity study. Investigators were not blinded to the group identity during experiments and data analysis except for the pathology assessments. Each *in vivo* experiment was performed once. Group sizes, replicate numbers, and statistical tests used are specified in corresponding result sections.

### Animal husbandry and ethics

All mouse immunogenicity experiments were approved by the local animal welfare committee (Landesuntersuchungsamt Rheinland-Pfalz, protocol G 20-12-005) and conducted at BioNTech SE according to the Federation of European Laboratory Animal Science Associations (FELASA) recommendations and in compliance with the German animal welfare act and Directive 2010/63/EU. Mice were kept under barrier and specific-pathogen-free (SPF) conditions in individually ventilated cages (maximum five animals of one sex per cage) and under controlled environmental conditions (20–24°C, 45–55% relative humidity, 75 AC/hr). Cages contained dust-free bedding made of debarked chopped aspen wood and additional nesting material. Autoclaved ssniff^®^ R/M-H food (ssniff Spezialdiäten GmbH) and autoclaved tap water were provided ad libitum and changed at least once weekly. Female C57BL/6JOlaHsd mice (Janvier labs) and female BALB/c mice (Janvier Labs) were used at 8 weeks of age. HLA-A2.1/DR1 transgenic mice were acquired from INFRAFRONTIER (ID: EM:01783) and bred in-house; male and female mice were used at 20–24 weeks of age.

For the GLP-compliant toxicity study, Wistar Hannover rats (Charles River) were housed in a limited access rodent facility at European Research Biology Center S.r.l. Italy (a fully accredited test facility by AAALAC) in clear polysulfone solid-bottomed cages (maximum five animals of one sex per cage) and under controlled environmental conditions (20–24°C, 40–70% relative humidity, 15–20 air circulation per hour (AC/hr). Nesting material was provided inside suitable bedding bags and changed at least twice per week. A commercially available laboratory rodent diet (4 RF 21, Mucedola S.r.l.) and drinking water were offered ad libitum throughout the study, except before urine collection. Procedures and facilities were compliant with the requirements of the Directive 2010/63/EU on the protection of animals used for scientific purposes. The national transposition of the Directive is defined in Decreto Legislativo 26/2014. Aspects of the protocol concerning animal welfare were approved by the local animal-welfare body (Organismo preposto al benessere animale). Rats were used at 8–9 weeks of age and allocated to groups by computerized stratified randomization to give approximately equal initial group mean body weights.

For the challenge studies, C57BL/6J mice (The Jackson Laboratory) were housed in SPF facilities at Trudeau Institute (Saranac Lake, New York, USA). Experimental mice were age and sex-matched. For the H37Rv study, mice were vaccinated at 12–13 weeks of age age. For the HN878 study mice were vaccinated at 8–9 weeks of age. All mice were treated in accordance with the National Institutes of Health guidelines for housing and care of laboratory animals and in accordance with the Institutional Animal Care and Use Committee guidelines at the Trudeau Institute under the approved protocols 13-003.12 and 13-003.14. All efforts were made to minimize suffering and pain as described in this approved protocol.

### Cell culture

HEK293T cells (ECACC #12022001) were grown in Dulbecco′s Modified Eagle′s Medium (DMEM, high glucose, GlutaMAX, pyruvate; Gibco) supplemented with 10% non-heat-inactivated fetal bovine serum (FBS) Superior (Sigma-Aldrich).

### RNA design and production

BNT164a1 and BNT164b1 contain four mRNAs where each mRNA encodes a fusion of two *Mtb* antigens: Ag85A-Hrp1, ESAT-6-RpfD, RpfA-HbhA and M72-VapB47. Antigen sequences were derived from H37Rv *Mtb* strain and exact amino acids included are listed in Table S1. A human HLA-A-derived signal peptide was added to the N-terminus of each fusion RNA to facilitate secretion^63^, and no linkers were introduced between antigens. M72 contained an N-terminal HIS-tag only in the construct used for the immunogenicity study in C57BL/6 mice and was subsequently removed.

BNT164a1 and BNT164b1 DNA templates were cloned into a plasmid vector with backbone sequence elements as described previously^16^. The DNA was purified and spectrophotometrically quantified. The four DNA templates were mixed and *in vitro*- transcribed by T7 RNA polymerase in the presence of a trinucleotide cap1 analogue ((m27,3′-O)Gppp(m2’-O)ApG) (TriLink). For BNT164b1, N1-methylpseudouridine-5′- triphosphate (m1ΨTP) (Thermo Fisher Scientific) was used instead of uridine-5′- triphosphate (UTP). RNA was purified using magnetic particles. RNA integrity was assessed by microfluidic capillary electrophoresis (Agilent Fragment Analyzer), and the concentration, pH, osmolality, endotoxin level and bioburden of the solution were determined.

### RNA formulation

Purified RNA was formulated into lipid nanoparticles as described before (Acuitas Therapeutics Inc)^16^. The lipid nanoparticle contains RNA, an ionizable lipid, ((4- hydroxybutyl)azanediyl)bis(hexane-6,1-diyl)bis(2-hexyldecanoate)), a PEGylated lipid, 2- [(polyethylene glycol)-2000]-N,N-ditetradecylacetamide and two structural lipids (1,2- distearoyl-sn-glycero-3-phosphocholine (DSPC]) and cholesterol).

### Cell transfection

HEK293T cells at 80–90% confluency were seeded in 12-well plates at 2 × 10^5^ cells/well one day prior to the transfection. HEK293T cells were transfected with uRNA or modRNA constructs (0.25 µg for single mRNAs, or 1 µg for mix of all four mRNAs) using RiboJuice^TM^ mRNA Transfection Kit (Sigma-Aldrich) according to the manufacturer’s instructions. Briefly, mRNA was diluted in OptiMEM and mixed with mRNA boost reagent and RiboJuice^TM^ mRNA reagent. The transfection mixture was incubated for 4 min at room temperature (RT) and added dropwise to the cells. The plate was gently mixed and incubated overnight (18 hours [h]) at 37°C, 5% carbon dioxide (CO_2_), in a humidified atmosphere. Non-treated control wells were run in parallel (non-transfected cells).

### Cell lysis and total protein isolation

Transfection media was discarded from transfected cells, which were rinsed with Dulbecco’s phosphate-buffered saline (DPBS) and centrifuged at 300 ×g for 5 minutes (min) at 4°C. The supernatant was discarded, 100 µL radioimmunoprecipitation assay (RIPA) buffer with 0.1% sodium dodecyl sulfate (SDS) containing a protease inhibitor cocktail (one cOmplete™ ULTRA) was added to the tube and incubated for 10 min on ice to lyse cells. The tube was centrifuged at ∼17,000 ×g for 10 min at RT and the supernatant containing the total protein suspension was kept on ice and used immediately to determine protein concentration for western blot, or stored at –80°C.

### Western blot

Total protein extracts from transfected and non-transfected control HEK293T cells were quantified with Pierce™ BCA protein assay kit (Thermo Scientific), according to the manufacturer’s protocol. Total protein extract and recombinant protein controls (produced in house in an *Escherichia coli* BL21 (DE3) expression system) were denatured under reducing conditions (95°C) and subjected to SDS-PAGE. 10 µg total protein per well or pre-stained protein marker (3 µL PageRuler™ Plus; Thermo Fisher Scientific) were loaded to the 8–16% polyacrylamide gradient gel (Mini PROTEAN® TGX™; Bio-Rad) and run at 140 V for ∼40 min. Proteins were transferred to a nitrocellulose membrane using a semi-dry method (Trans-Blot Turbo Transfer System, Bio-Rad). Blotted proteins were detected with primary antibodies (mouse anti-ESAT6, Abcam #ab26246; rabbit anti-Hrp1, BEI # NR-36512; mouse anti-RpfD, anti-RpfA, anti-HbhA, anti-M72, anti-VapB47, anti-Ag85A, all made-to-order Squarix GmbH; rabbit anti-tubulin, Cell Signaling Technology #9099S) and secondary antibodies (anti-mouse IgG, Jackson ImmunoResearch Laboratories # 115-035-071; anti-rabbit IgG, Sigma-Aldrich #A0545). Blots were developed with chemiluminescent substrate (Cytiva Amersham™ ECL™ Prime Western Blotting Detection Reagent) and analyzed with the ChemiDoc Imaging System and ImageLab software (Bio-Rad).

### Immunizations

For immunization studies, the mice were anesthetized by inhalation of 2.5% isoflurane in oxygen and the injection site on the hind leg of the mouse was shaved. The mRNA test items at a dose of 4 µg or saline were injected IM into the gastrocnemius muscle in a volume of 20 µL per injection with a 30g needle insulin syringe. BCG (10^6^ CFUs) was administered SC without prior anesthesia. Injection sites were observed every 24 h for 2 days post injection and at least once weekly throughout the study.

For the toxicity study, the rats received IM injections of the mRNA test items or saline into the quadriceps muscle (alternating the right and left thigh) with a 30 gauge (g) needle insulin syringe, at the dose volume as indicated: 60 µL saline, 30µg/60 µL NTL, 10µg/20 µL or 30µg/60 µL BNT164b1, or 10µg/20 µL BNT164a1. Prior to the first administration, and as necessary during the course of the study, the thigh was shaved. A daily examination of the injection site was performed at pre-dose, and approximately 4 and 24 hours after each administration.

For the challenge study, the mice were immunized IM in the gastrocnemius muscle with 4 µg of mRNA test items or saline in a volume of 20 µL per injection using a 30g needle. For SC immunization, a 26g needle was used and BCG-containing saline (10^6^ CFU) was injected either at the nape of the neck or around the hip. Injection sites were monitored weekly or on consecutive days following an injection.

### Animal monitoring

Routine animal monitoring was carried out daily including inspection for dead mice and control of food and water supplies. The health of each mouse was closely assessed at least once weekly, and the results were documented in health monitoring sheets. The general physical condition of the mice was assessed according to the following parameters: mice were observed daily and weighed at least weekly for the duration of the study. Animals that lost more than 20% body weight were euthanized immediately. Mice were observed for the following clinical signs for the duration of the study: 1) moderate toxicity signs: intermittent hunching, ruffled coat, intermittent abnormal breathing, or intermittent tremors and convulsions; 2) substantial toxicity signs: prolonged immobility, hunched, dehydration, skin tenting, labored breathing, or prolonged prostration. Animals exhibiting signs of substantial toxicity were euthanized immediately. During the immunization study in HLA-A2.1/DR1 transgenic mice, one animal in the BNT164a1 group was euthanized due to treatment-unrelated health issues and could not be included in the analysis. During the *Mtb* H37Rv challenge study, one animal in the BNT164a1 group was euthanized due to severe alopecia and erythema (>2.0 cm) at the injection site after the second immunization, and one animal in BNT164b1 prime-boost group was euthanized due to an unrelated event. In the *Mtb* HN878 challenge study, one mouse immunized with BCG had to be sacrificed with symptoms of a possible ear infection.

In the toxicity study, rats were checked at least once daily for mortality and clinical signs. Each animal was weighed on the day of allocation to vaccination group, on the day of administration, 24 and 48 hours after each administration, and just prior to necropsy. During recovery, animals were weighed at weekly interval. Water consumption for each cage was recorded daily, and food consumption was recorded twice weekly. Rectal body temperature was measured on Days 1, 8, 15, 22 pre-dose and at approximately 4 h and 24 h post-dose (Days 2, 9, 16 and 23). In case of recorded post-dose rectal body temperature ≥ 38.5°C, the temperature was measured the day after (48 h post-dose) until the values returned below 38.5°C.

### Blood collection and serum isolation

In the immunization studies, blood was sampled either from the facial vein using a lancet (without prior anesthesia), or from the retro-orbital venous sinus using a 29 g glass capillary (upon isoflurane-induced anesthesia). Blood was collected into an appropriate plastic tube (Sarstedt, Z-gel included for clotting activation). Blood samples were centrifuged for 5 min at RT and 10,000 ×g to collect serum, which was then stored at −20°C.

In the toxicity study in rats, blood samples for clinical pathology investigations were collected under condition of overnight food deprivation from the retro-orbital sinus on day 3 or from the abdominal vena cava on days 24 and 43. Blood samples were collected into tubes as follows: EDTA anticoagulant for hematological investigations; citrate anticoagulant for coagulation tests (fibrinogen); without anticoagulant for biochemical tests, alpha 2- macroglobulin, and alpha 1-acid-glycoprotein.

### Euthanasia procedure and organ collection

Mice were euthanized by cervical dislocation or by exposure to CO_2_.

Rats were euthanized by exsanguination under isoflurane anesthesia and necropsy was supervised by a pathologist. A detailed *post mortem* examination was conducted, observations were noted, the requisite organs weighed and the required tissue samples preserved in fixative (10% neutral buffered formalin) and processed for histopathological examination.

### Splenocyte preparation

Spleens were collected and stored in DPBS on ice. Single-cell suspensions were prepared by homogenizing through 70 µm cell meshes using the plunger of a syringe. Splenocytes were washed with an excess volume of DPBS followed by centrifugation at 300 ×g for 6 min at RT and discarding the supernatants. Erythrocytes were lysed with erythrocyte lysis buffer (154 mM NH_4_Cl, 10 mM KHCO_3_, 0.1 mM EDTA) for 5 min at RT. The reaction was stopped with an excess volume DPBS. After another washing step, cells were resuspended in complete RPMI medium (10% FBS, 1% NEAA, 1% sodium pyruvate, 0.5% penicillin/streptomycin), passed through a 70 µm cell mesh again, counted and stored short-term at 4°C for use on the same day or frozen in Cryomedium (FBS non heat-inactivated + 10% DMSO cell culture grade).

### Sorting of splenocytes

CD4^+^ and CD8^+^ T cells were sorted from single-cell suspensions of splenocytes by magnetic-activated cell sorting (MACS®, Miltenyi Biotec). Prior to cell sorting, splenocytes were pooled from all five mice in each group. Mouse CD4^+^ and CD8^+^ T-cell isolation kits, QuadroMACS™ Separator, and LS columns were used, following the manufacturer’s instructions. Negative isolation method was used, which resulted in elution of untouched CD4^+^ and CD8^+^ T cells. Cells were separately centrifuged and prepared at a concentration of 1×10^6^ cells/mL in RPMI 1640 medium supplemented with 10% heat-inactivated FBS, 1% NEAA, 1% sodium pyruvate, 1% HEPES, 0.5% penicillin/streptomycin, 50 µM 2- mercapthoethanol.

### Preparation of bone marrow-derived dendritic cells (BMDCs)

For analyzing the sorted CD4^+^ and CD8^+^ T cells via ELISpot, BMDCs served as antigen-presenting cells. BMDCs were differentiated from the corresponding mouse model by stimulating bone marrow cells with GM-CSF (1000 U/mL) for 7 days. Dendritic cell identity was confirmed by expression of CD11c, CD86, and MHC-II via flow cytometry.

### ELISpot

ELISpot assays were performed according to the manufacturer’s protocol (with minor modifications as described below) using the mouse IFN-γ ELISpot^PLUS^ kit (Mabtech). Briefly, 96-well ELISpot plates pre-coated with IFN-γ antibody were washed with PBS and blocked with serum-free medium for at least 30 min at RT. The medium was removed, and cells were added to the plates: either 5×10^5^ total splenocytes/well (100 µL), or 1×10^5^ sorted CD4^+^/CD8^+^ T cells/well (100 µL) together with 5 x 10^4^ BMDCs/well (50 µL). BMDCs served as antigen-presenting cells for sorted T cells. Another 100 µL of overlapping peptide pools (15-mer long peptides with 11 amino acid overlap) were added to the wells at 2 µg/mL final concentration: Ag85A (34 overlapping peptides; Pepscan), M72 (180 overlapping peptides; Pepscan), ESAT6 (21 overlapping peptides; Pepscan), HbhA (47 overlapping; Pepscan), RpfD (36 overlapping peptides; Pepscan), Hrp1 (33 overlapping peptides; JPT Peptide Technologies), RpfA (99 overlapping peptides; Pepscan), VapB47 (22 overlapping peptides; Pepscan). PPD (AJ vaccines) was used at 10 µg/mL. As a positive control the splenocytes were stimulated with 2 µg/mL concanavalin A (Sigma), and for an unstimulated control only medium was added. TRP1 (for C57BL/6JOlaHsd mice), AH1 (for BALB/c mice) or CMVpp65 (for HLA-A2.1/DR1 transgenic mice) (all JPT Peptide Technologies) were used as non-specific stimulants at 2 µg/mL. Plates were incubated overnight in a humidified incubator with 5% CO_2_ at 37°C for 18 h. Next, the detection antibody, Streptavidin-ALP, and the ready-to-use substrate were added to the wells according to the manufacturer’s protocol. After plate drying for 2–3 h under the laminar flow or overnight, an ELISpot plate reader (ImmunoSpot S6 Core Analyzer, CTL) was used to count and analyze spot numbers per well.

### T-cell phenotyping of non-infected animals via flow cytometry

5×10^5^ fresh splenocytes in 100 µL RPMI 1640 medium supplemented with 10% heat-inactivated FBS, 1% NEAA, 1% sodium pyruvate, 1% HEPES, 0.5% penicillin/streptomycin, 50 µM 2-mercapthoethanol were transferred to 96-well U-bottom cell culture plates. 100 µL of an overlapping peptide mix containing all eight *Mtb* antigens (as specified above) were added at 1 μg/mL per peptide in the presence of co-stimulatory antibodies (1 μg/mL anti-CD28 (eBioscience, clone 37.51) and 1 μg/mL CD49d (Biolegend, clone R1-2)). As a non-stimulation control, only medium was added to detect unspecific background signals. Plates were incubated for 1 h in a 37°C humidified incubator with 5% CO_2_ before adding GolgiStop^TM^ + GolgiPlug^TM^ (BD Biosciences). After another 4–5 h incubation, the plates were stored at 4°C overnight. Cells were then harvested, transferred to a 96-well, U-bottom plate for flow cytometry, centrifuged (360xg, 5 min, 4°C), resuspended in 100 µL per well 1x Permeabilization buffer (eBiosciences), and incubated for 5 min at 4°C. After another centrifugation (400xg, 3 min, 4°C), the supernatant was discarded and the cells were stained for 20 min at 4°C using the following panel: Live/Dead-APC-R700 (1:1000, BD Biosciences, #564997), CD3-BV605 (1:200, Biolegend #100351), CD4-BV510 (1:400, Biolegend #100559), CD8-APC (1:200, BD Biosciences #553035), IFNγ-PE-Cy7 (1:00, Biolegend #505826), IL2-BV711 (1:100, Biolegend #503837), TNFα-Alexa488 (1:100, Biolegend #506313), CD127-PE (1:200, Biolegend #121111), KLRG1-APC_eFluor780 (1:100, Thermo Fisher, #47-5893-82), and CD62L-BV785 (1:200, Biolegend #104440). After the staining procedure, cells were washed twice in Permeabilization buffer, and resuspended in 150 µL FACS buffer (PBS +0.1% BSA) for flow cytometry analysis using FACSCelesta (BD).

### T-cell phenotyping via flow cytometry of *Mtb* HN878-infected animals

Lung tissue was prepared by coarsely chopping and enzymatically digesting the tissue in a 0.5 mg/mL solution of collagenase (Roche) and DNase (Sigma Aldrich) and incubated for 30–45 minutes at 37°C. Single cell suspensions were prepared from digested lung tissue or mesenteric lymph nodes harvested into HBSS by dispersing through a 70-μm nylon tissue strainer (BD Falcon). The cell suspensions were treated with Gey’s solution to remove erythrocytes. Lymphocytes were enriched from lung tissue by differential centrifugation, using a gradient of 40/80% Percoll (GE Healthcare).

Lymphocytes were transferred to a 96-well plate, at a concentration of 1.5 to 2 x 10^6^ cells per well. After incubating the lymphocytes with Fc block (Biolegend) for 15 minutes on ice, they were stained with tetramer reagents (PE-labeled ESAT-6_1-20_ I-A^b^ Tetramer (MBL Life science, product code MBL-TS-M707-1) and APC-labeled Mtb32A(PepA)_309-318_ H-2D^b^-Tetramer (produced at Trudeau institute) for 60 minutes at room temperature, and a surface antibody panel for 30 minutes on ice. Surface antibody panel consisted of: CXCR5-FITC (clone 7E.17G9, Biolegend #145520); ; CD127-PE-CF594 (clone SB/199, BD #562419); KLRG1-PerCP-Cy5.5 (clone 2F1/KLRG1, BD #563595); PD1-Pe-Cy7 (clone RMP1-30, Bioloegend #109110); CD4-BV421 (clone RM4-5, Biolegend #100544); CD8-BV510 (clone 53-6.7, Biolegend #100751); CD62L-AF700 (clone MEL-14; Biolegend #104426); CD44.APC-eFluor780 (clone IM7; Invitrogen #47-0441-82) and Zombie UV Live/Dead stain (BioLegend #77474). Stained cells were fixed in 2% paraformaldehyde for 24 h before being analyzed on a LSRII flow cytometer (BD Biosciences). Flow cytometry data were analyzed using FlowJo software version 10.8.1.

### ELISA

Serum samples were tested in 96-well plates for their antigen-specific antibody concentration. Each well of a MaxiSorp^TM^ plate was coated with 100 ng/100 µL per well recombinant *Mtb* proteins produced in *E. coli* or mouse IgG isotype control. For the lysate ELISA 1 µg/100 µL of *Mtb* H37Rv lysate (BEI resources NR-14822) was used instead. Plates were incubated at 4°C overnight. Wells were washed with PBS with 0.01% Tween 20 (PBS-T) and blocked with casein blocking buffer (Sigma-Aldrich) at 37°C for 1 h on a shaker. After another round of washing, serially diluted serum samples in blocking buffer or negative control (blocking buffer alone) were added in duplicate and incubated at 37°C for 1 h on a shaker. After washing the plates, HRP-conjugated goat anti-mouse IgG secondary antibody was added (Jackson ImmunoResearch #115-035-071, 1:7500 in blocking buffer), and plates were incubated at 37°C for 45 min on shaker. The plates were washed again, TMB substrate (Biotrend Chemikalien GmbH) was added and incubated for 8 min at RT in the dark. The reaction was stopped with 100 µL 25% sulfuric acid and the absorbance (450 nm, ref. 620 nm) was measured on Epoch microplate reader (BioTek). The mean changes in optical density (ΔOD) measured from the absorbance values at 450 nm and 620 nm for each sample were calculated. IgG concentrations were quantified by interpolation from the IgG isotype standard curve in GraphPad Prism 9 or Genedata Screener.

### Toxicity study blood analysis

Hematological parameters (e.g., white blood cells and large unstained cells) were measured using Siemens Advia 120. Fibrinogen values were determined by Instrumentation Laboratory ACL Elite PRO. Alpha-2-macroglobulin and alpha 1-acid-glycoprotein were analyzed by validated commercial ELISA kits. Some samples were unsuitable for haematological analysis: day 3: 2 animals from saline group, BNT164 NTL group, and BNT164b1 30 μg/mL group each; day 24: 2 animals from saline group, 1 animal from BNT164 NTL group, 1 animal from BNT164b1 10 μg/mL group; day 43: 1 animal from saline group.

### Challenge studies

C57BL/6J mice between 12 and 13 weeks old were immunized IM with mRNA test items as follows: BNT164a1, BNT164b1 and NTL were given twice (days 0 and 21). A comparison group was given 10^6^ CFUs of BCG SC once (day 0). On day 56, animals were challenged with a low dose (∼100 CFU) aerosol *Mtb* (strain H37Rv or HN878), using a Glas-Col airborne infection system. Stocks of *Mtb* strains used in the aerosol infection were cultured in Proskauer Beck medium containing 0.05% Tween 80 to mid-log phase and frozen in 1 mL aliquots at –80°C. Animals were euthanized either 30 or 60 days post-challenge (Days 87 and 117 post-vaccination), left lung lobes and spleens were harvested and homogenized in 4.5 mL of normal saline using glass tube and pestles (Glas-Col). Organ homogenates were serially diluted 1:10 and 100 µL of each dilution step were plated on 7H11 agar on a 100 mm petri dish. Agar plates were incubated for 18 days at 37°C, 6% CO_2_ in 60% humidity. After the indicated incubation time, bacterial colonies were counted and the total bacterial burden in tissue homogenate was calculated and recorded as log_10_ CFU.

### Histopathological examinations

For the rat samples, fixed and preserved tissues were dehydrated, embedded in paraffin wax, cut in 5 µm thick sections, and stained with hematoxylin and eosin.

For the murine samples, right lung lobes were inflated with 0.5-0.7 mL neutral buffered formalin, placed into histology cassettes (Fisher Scientific #15-182-706), and submerged in 10% neutral buffered formalin for 48 hours. The fixed tissues were processed using Tissue-Tek VIP 3000 (Sakura) and embedded within paraffin wax (Fisher Scientific 22900701) using a Tissue-Tek Embedding station. Once set, the blocks were sectioned at 5 microns using a Leica RM2125RT microtome and placed on slides (Superfrost PLUS; Fisher Scientific 1255015). Once dried, a representative slide from each mouse was heated for 20 minutes on a slide warming plate to adhere the tissues to the slides. After deparaffinization and rehydration, the slides were stained with hematoxyolin (Epredia) and eosin (Epredia) or submitted to acid-fast staining using the Acid-Fast Stain Kit (ENG Scientific 6500) following the manufacturer’s protocol. The slides were cover slipped using EcoMount (BioCare Medical, EM897L) and 22mmX22mm coverslips (Fisher Scientific, 12541016). Full slide scans were taken with the 40x objective using a Leica Aperio AT2 Slide Scanner. Granuloma burden per lung section was quantified by a pathologist blinded to the group identities using the following scoring system: Grade 0: no granulomas observed; Grade 1: Minimal, <5 granulomas; Grade 2: Mild, 5–9 granulomas; Grade 3: Moderate, 10–14 granulomas; Grade 4: Marked, 15–20 granulomas; Grade 5: Severe, >20 granulomas. The cellular composition of granulomas was further classified by assigning a rounded percentage of the infiltrate/aggregates contributed by macrophages, lymphocytes, or neutrophils (0, 25, 50, 75 or 100%); these percentages were then multiplied by the granuloma score to obtain an individual score for each cell type. Using acid-fast-stained slides, the number of discrete granulomas containing acid-fast, rod-shaped bacteria, consistent with *Mtb* morphology, were enumerated. The percentage of granulomas containing acid-fast bacteria was calculated based on the granuloma count (above).

### Statistical analysis

GraphPad Prism 9 software was used for generating graphs and for performing statistical analysis. Flow cytometry and ELISpot (complete splenocytes) data were analyzed by 1-way ANOVA with Tukey’s multiple comparisons test. *Mtb* lysate ELISA data were analyzed by ratio-paired t tests corrected for multiple comparisons by false discovery rate calculation using Benjamini, Krieger, Yekutieli with a threshold of q < 0.05. Colony counts and bacterial burden assessment via pathology in challenge studies were analyzed by 1-way ANOVA with Dunnett’s multiple comparison test. Immune cell quantification and phenotyping data in challenge studies were analyzed by Kruskal-Wallis ANOVA followed by Dunńs multiple comparison test. An alpha significance level of 0.05 was set as threshold for p value (95% confidence interval). Statistical significance was reported as *p < 0.05, **p < 0.01, ***p < 0.001, ****p < 0.0001.

## DATA AVAILABILITY

All data associated with this study can be found in the main text or the Supplementary Information. Source data are available from the corresponding author upon reasonable request.

## ACKNOWLEDGEMENTS

This work was supported, in whole or in part, by the Gates Foundation. Under the grant conditions of the Foundation, a Creative Commons Attribution 4.0 Generic License has already been assigned to the Author Accepted Manuscript version that might arise from this submission. We thank Sonja Witzel, from TRON gGmbH, for assistance with molecular cloning; Thomas Hiller for production of recombinant proteins; Thomas Ziegenhals and team for RNA development and BioNTech Formulation & Drug Delivery Unit for formulation services; BioNTech Animal Facility (especially Phillip Windecker and Henrike Fleige) for performing and supervising *in vivo* immunogenicity experiments; Julia Haupt for leading project management; Robbert van der Most and Kellen Faé for valuable input and fruitful discussions; Katja Schlatterer and Theo Benjamin Stahl for assay development; William Reiley from TICRO BioServices for his valuable input in the challenge study design. The following reagents were obtained through BEI Resources, NIAID, NIH: Mycobacterium tuberculosis, Strain H37Rv, Whole Cell Lysate, NR-14822 and polyclonal anti-*Mycobacterium tuberculosis* HRP1 (Gene Rv2626c) (antiserum, Rabbit), NR-36512. Figures 1A, 2A, 4A, and 5A were created with BioRender.com

## Contributions

A.B.V. and U.Ş conceived the study. N.A., L.S.A., S.S., A.C., A.B.V., J.D., and M.D. designed the experiments. N.A., S.S., and J.V. performed experiments. N.A., L.S.A., S.S., A.C., J.V. and N.V. analyzed the data. N.A., L.S.A., S.S., A.C., N.V., A.B.V., J.D., M.D., and U.Ş. interpreted the data. L.S.A. and N.V. drafted the manuscript. All authors were involved in reviewing and editing the manuscript.

