## Supplementary information for "mRNA-based tuberculosis vaccines BNT164a1 and BNT164b1 are immunogenic, well-tolerated and efficacious in rodent models"

### Table S1. *Mtb* antigens included in BNT164 vaccine candidates

| Antigen* | Characteristics* | AAs in BNT164 | Immunogenicity and efficacy |
|---|---|---|---|
| Ag85A (Rv3804c) | Has mycolyltransferase activity; essential for cell wall integrity; secreted protein. | 42–338 | Human: Humoral, $CD4^+$ and $CD8^+$ T-cell responses after boost immunization of healthy BCG vaccinated adults [46].<br>Mouse: Immunogenicity and protection against TB reported from DNA vaccine [47]. **NHP: Multiantigen cytomegalovirus-based vaccine was highly efficacious and even resulted in sterilizing immunity in a subgroup of animals [27]. |
| ESAT-6 (Rv3875) | Essential *Mtb* virulence factor; part of RD1 loci of *Mtb*; abundantly secreted at early stages of infection; secreted protein. | 2–95 | Human: CAF01-adjuvanted Ag85B/ESAT-6 fusion protein vaccination induced long-lasting $CD4^+$ T-cell response [48]. ESAT6 is one of the components of IGRA test used to diagnose latently TB infected individuals.<br>Mouse: Immunogenicity and protection against TB reported [49]. Deletion of ESAT-6 and CFP10 from MTBVAC (*Mtb* live-attenuated vaccine candidate) reduced the protection of MTBVAC [50]. **NHP: see above. |
| M72 (Rv0125 and Rv1196) | Fusion protein consisting of two antigens [51]:<br>Mtb32A (PepA): probable serine protease involved in metabolism and respiration; detected in early stage of infection; secreted protein.<br>Mtb39a (PPE18): PE/PPE family protein of unknown function; detected in late stages of infection; probable membrane-bound protein. | Similarly fused as described before [52] | Human: Immunization led to early onset humoral response and $CD4^+$ T-cell responses, sustained for 3 years; 49.7% (95% CI: 2.1 to 74.2) vaccine efficacy observed 3 years after immunization in phase II trial [28].<br>Mouse: Immunogenicity and protection against TB reported [51]. |
| VapB47 (Rv3407) | Possible antitoxin; virulence associated; intracellular protein. | 2–99 | Human: VapB47-specific IFNγ responses have been observed upon restimulating active TB patient blood cells [53].<br>Mouse: Immunogenicity and protection against TB reported when delivered as a DNA vaccine booster to BCG immunization [54]. **NHP: see above. |
| Hrp1 (Rv2626) | Dormancy associated; expression induced by oxygen/nutrient stress; detected in culture as well as membrane-bound. | 2–143 | Human: Hrp1-specific antibody detected in active TB patients [55].<br>Mouse: Immunogenicity and protection against TB reported. Plasmid DNA vaccination induced strong humoral and Th1-type cellular response [56]. Long-term protection from TB when given together with ESAT6-Ag85B-$MPT64_{(190-198)}$-Mtb8.4 [57]. **NHP: see above. |
| RpfD (Rv2389c) | Probable role in promoting the resuscitation. | 2–154 | Human: $CD4^+$ and $CD8^+$ T-cell responses observed in LTBI and active TB patients [58]. IFNγ production by T cells in response to RpfD and RpfA is higher in LTBI than in pulmonary TB [59].<br>Mouse: Immunogenicity observed after *Mtb* infection and BCG vaccination. Moderate protection from TB was induced by immunization with a family member (RpfB) [60]. **NHP: see above. |
| RpfA (Rv0867c) | Probable role in promoting the resuscitation; detected in late stage of infection; secreted protein. | 2–407 | Similar to RpfD. |
| HbhA (Rv0475) | Required for extrapulmonary dissemination of *Mtb*; membrane-bound. | 2–199 | Human: T-cell response against HbhA reported from LTBI[61].<br>Mouse: Immunogenicity and protection against TB reported [62]. |

*Data source: https://mycobrowser.epfl.ch/

BCG = Bacillus Calmette-Guérin; IFNγ = interferon gamma; IGRA = IFN Gamma Release Assay; LTBI = latent tuberculosis infection; *Mtb* = *Mycobacterium tuberculosis*; NHP = non-human primate; Tb = tuberculosis

#### Supplementary figure 1

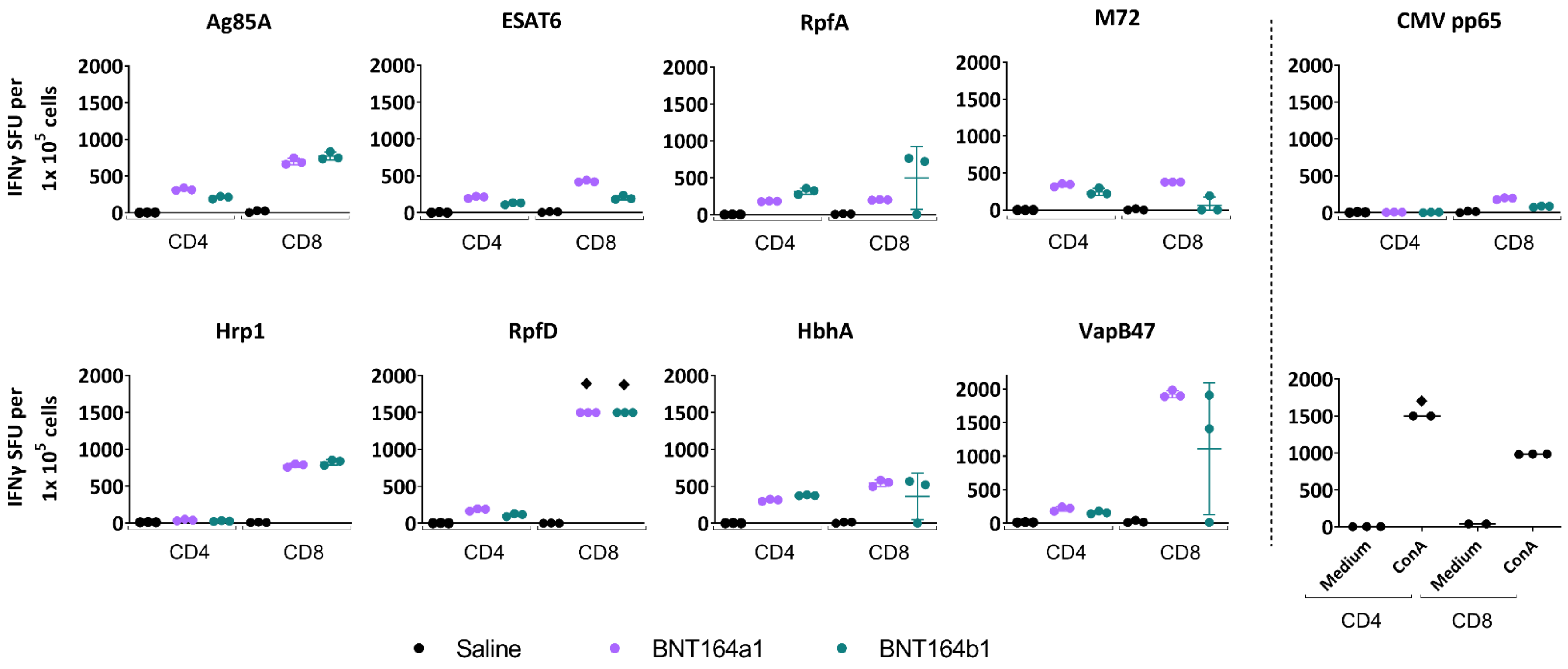


**Fig. S1. BNT164 candidates induced antigen-specific CD4⁺ and/or CD8⁺ T-cell responses in HLA-A2.1/DR1 humanized mice**

Splenocytes were isolated on Day 42 from mice injected twice (Days 0 and 21) IM with 4 µg of BNT164a1, 4 µg of BNT164b1, or a saline control. $CD4^+$ and $CD8^+$ T cells were magnetically sorted from splenocytes pooled from each treatment group (n=5/group) and stimulated with individual *Mtb* antigens, medium only, concanavalin A (con A), or non-specific peptide (CMVpp65). Bone marrow-derived dendritic cells from non-vaccinated mice (unrelated cohort of naïve mice) were co-cultured with T cells as antigen-presenting cells. The responses were assessed by IFNγ ELISpot assay after ~18 hours incubation. Circles represent duplicate/triplicate measurements; horizontal bars represent group means ± standard deviation. For some conditions, the upper limit in number of spots that can be correctly counted was reached (indicated with the rhombus symbol). SFU = spot-forming unit. IM = intramuscular.

### Supplementary figure 2

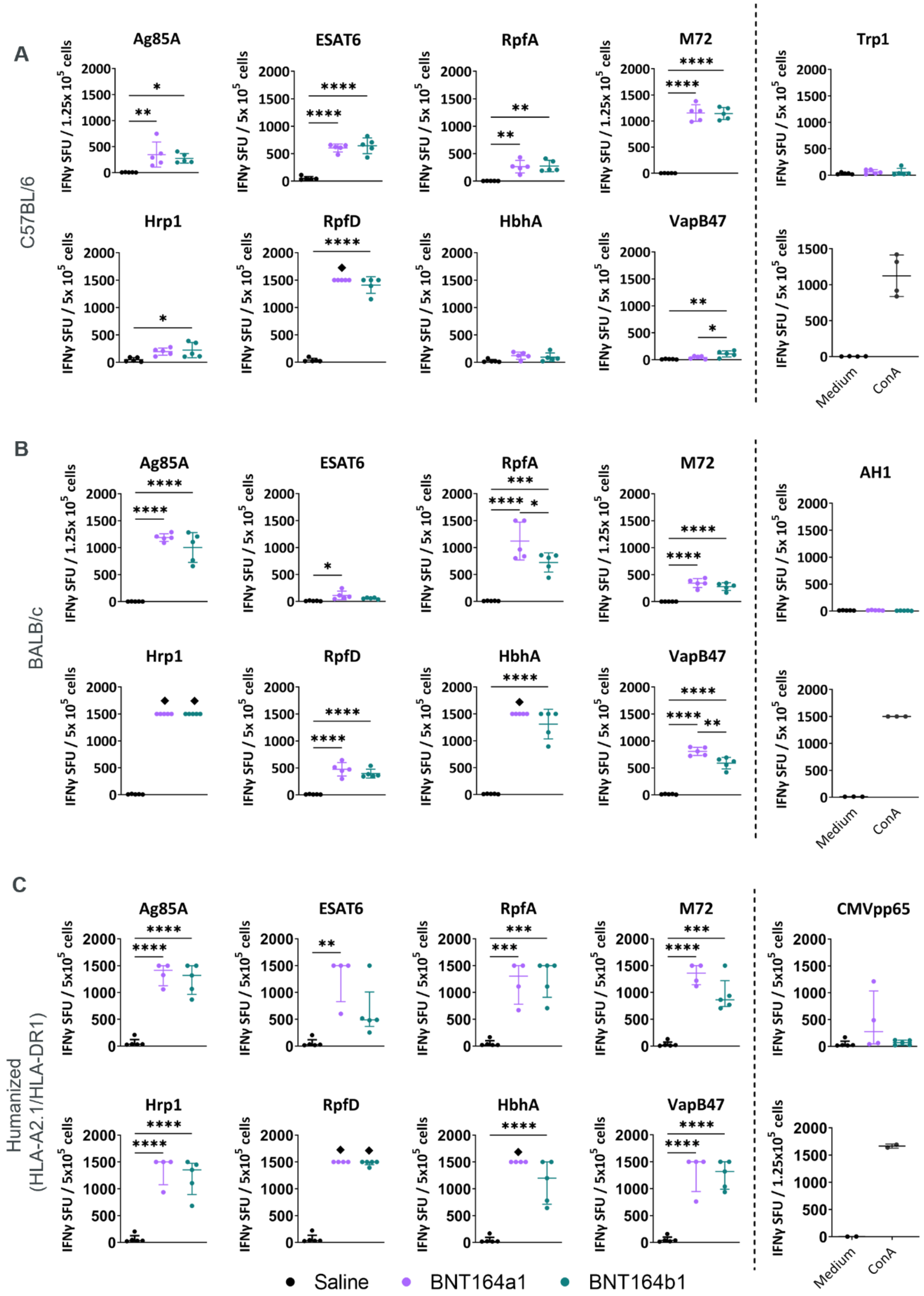

**Fig. S2. BNT164 candidates induced antigen-specific total T-cell responses in different mouse strains**
**(A)** C57BL/6, **(B)** BALB/c, and **(C)** HLA-A2.1/DR1 mice received IM injection of 4 µg BNT164a1, 4 µg BNT164b1, or saline control on Days 0 and 21. Splenocytes were isolated on Day 42 and stimulated with antigen-specific peptide pools, medium only, concanavalin A (con A), or non-specific peptides (TRP1, AH1, CMVpp65). Responses were assessed by IFNγ ELISpot assay after ~18 hours incubation. Group mean values are indicated by horizontal bars (± standard deviations), means from individual mice (n=4-5/group, measured in duplicates) are depicted as circles. For stimulation with medium or concanavalin A, technical replicates are shown. One-way ANOVA with Tukey's multiple comparisons test was performed**;** * = $p < 0.05$; ** = $p < 0.01$; *** = $p < 0.001$; **** = $p < 0.0001$. SFUs for some samples reached the upper limit in number of spots that can be correctly counted and were not statistically analyzed (indicated with rhombus symbol). IM = intramuscular; SFU = spot-forming unit.

### Supplementary figure 3

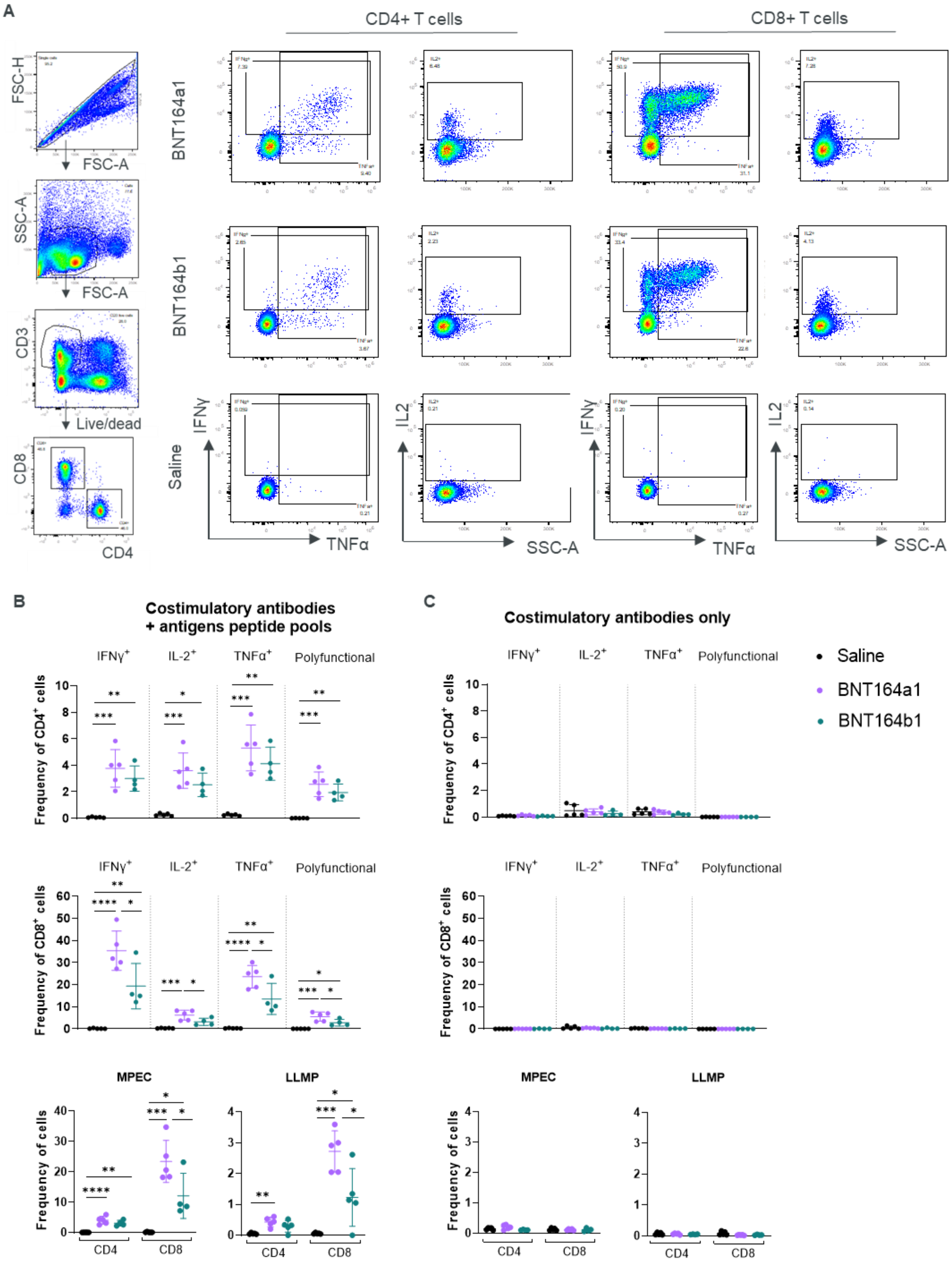

**Fig. S3. BNT164 candidates induced polyfunctional T cells and T cell memory**

Splenocytes were isolated on Day 42 from C57BL/6 mice injected twice (Days 0 and 21) IM with 4 µg BNT164a1, 4 µg BNT164b1, or saline control. Splenocytes were stimulated with costimulatory antibodies (anti-CD49d and anti-CD28) with or without a mix of overlapping peptide pools covering all the encoded antigens. Cells were stained for intracellular and extracellular markers including viability, CD3, CD4, CD8, IFNγ, IL-2, TNFα, CD127, KLRG1, and CD62L. (A) The gating strategy and representative flow cytometry dot plots. (B–C) Percentage of CD4/CD8 cells positive for single cytokines and polyfunctional T cells ($IFN\gamma^{+}$/$IL\text{-}2^{+}$/$TNF\alpha^{+}$) (top and middle) and percentage of MPEC ($TNFa^{+}$ or $IFN\gamma^{+}$/ $CD127^{+}$/$KLRG1^{-}$/$CD62L^{-}$) and LLMP ($TNFa^{+}$ or $IFN\gamma^{+}$/ $CD127^{+}$/$KLRG1^{-}$/$CD62L^{+}$) CD4/CD8 cells (bottom). Group mean values are indicated by horizontal bars (± standard deviations); individual mouse values (n=5/group) are depicted as circles (one sample from BNT164b1 group was excluded due to insufficient cell number). One-way ANOVA with Tukey's multiple comparisons test was performed: * = $p < 0.05$; ** = $p < 0.01$; *** = $p < 0.001$; **** = $p < 0.0001$.
IM = intramuscular; MPEC = memory precursor effector cell; LLMP = long-lived memory precursor cell.

### Supplementary figure 4

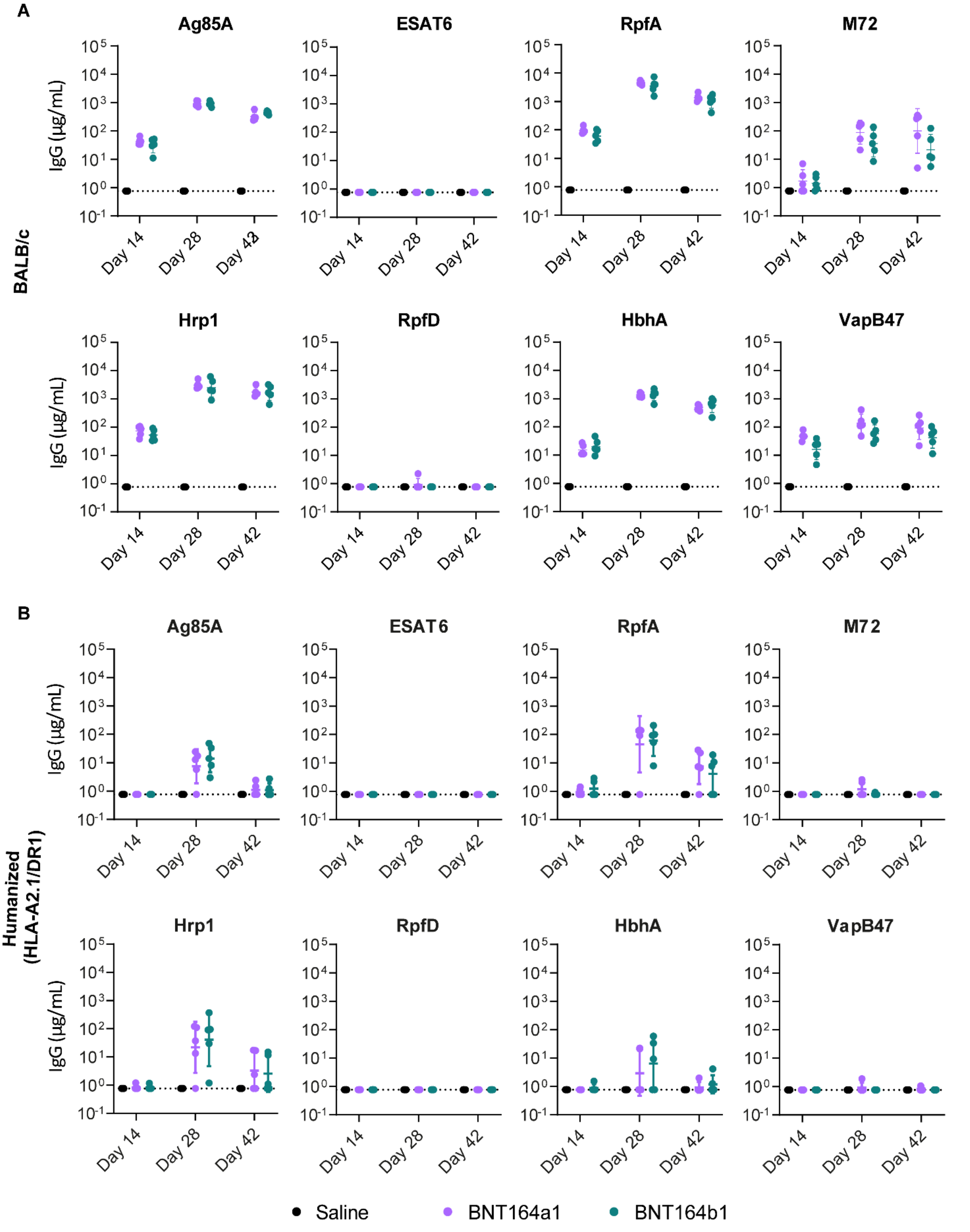


**Fig. S4. IgG responses in BALB/c and HLA-A2.1/DR1 humanized mice following prime-boost immunization with BNT164 candidates**

Mice were injected IM with 4 µg BNT164a1, 4 µg BNT164b1, or saline control on Days 0 and 21. Antigen-specific IgG antibodies were assessed by ELISA from sera isolated on indicated days. Group mean values are indicated by horizontal

bars (± standard deviations), means from individual mice (n=5/group, measured in duplicates) are depicted as circles. Dashed line indicates lower limit of detection (LLOD). In (B) one mouse in BNT164a1 group had to be sacrificed before the end of the experiment due to treatment-unrelated health issues and could not be included in the analysis.
IM = intramuscular.

#### Supplementary figure 5

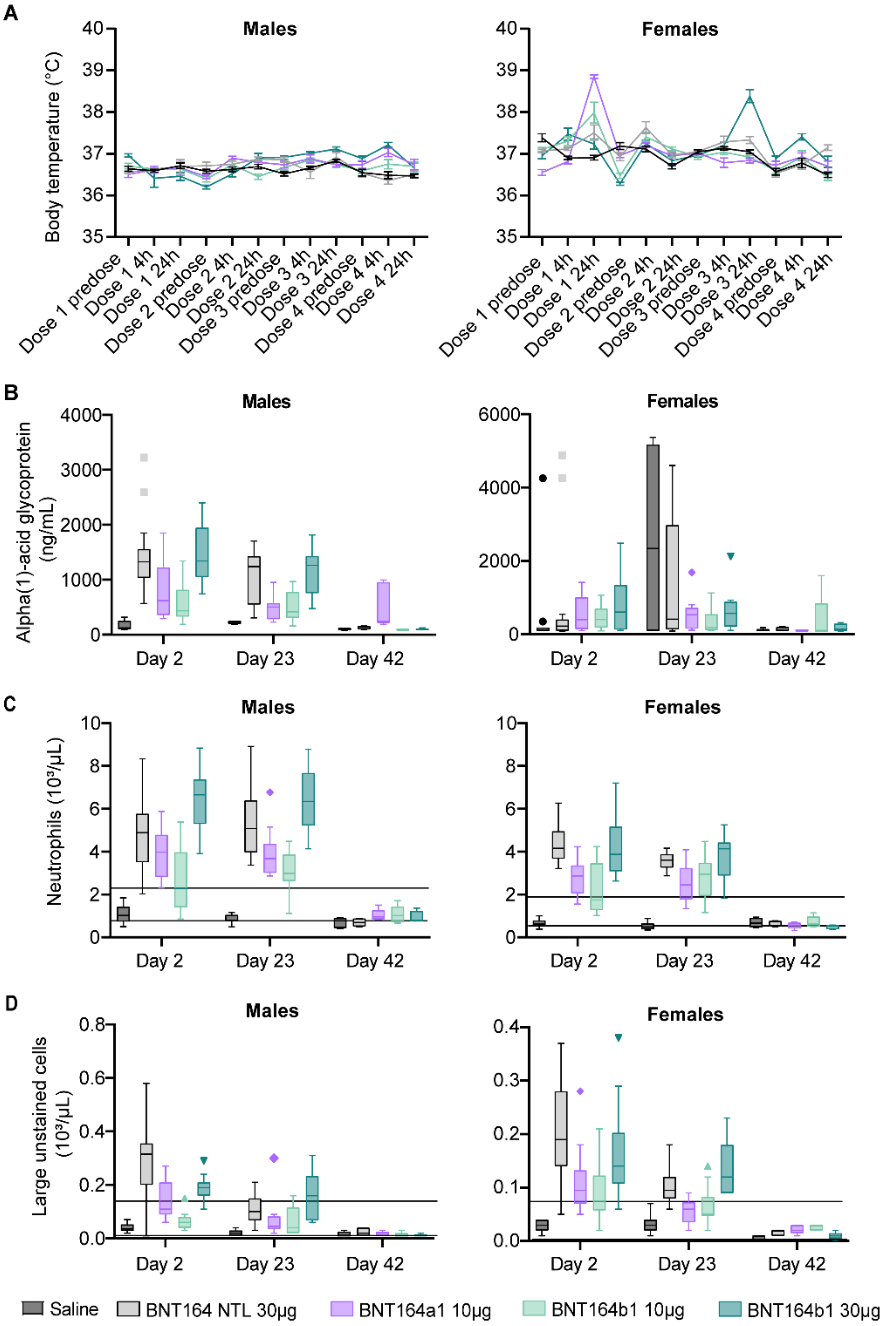


**Fig. S5. Additional readouts from a repeat-dose toxicity study in Wistar Han rats**

Wistar Han rats were injected IM with saline, BNT164 non-translatable control (NTL, 30µg), BNT164a1 (10µg), or BNT164b1 (10µg or 30µg), on Days 0, 7, 14, and 21. Main study group was sacrificed on Day 23 (n=10), while the recovery group was sacrificed on Day 42 (n=5). Some samples were unsuitable for hematological analysis following coagulation and/or following unreliable instrumental data and are indicated in the method section. (A) Animal rectal temperature (mean ± standard error of the mean). (B-D) Laboratory findings plotted with Tukey box plots. Horizontal lines in (C) and (D) represent historical values range for this strain at study site. IM = intramuscular.

#### Supplementary figure 6

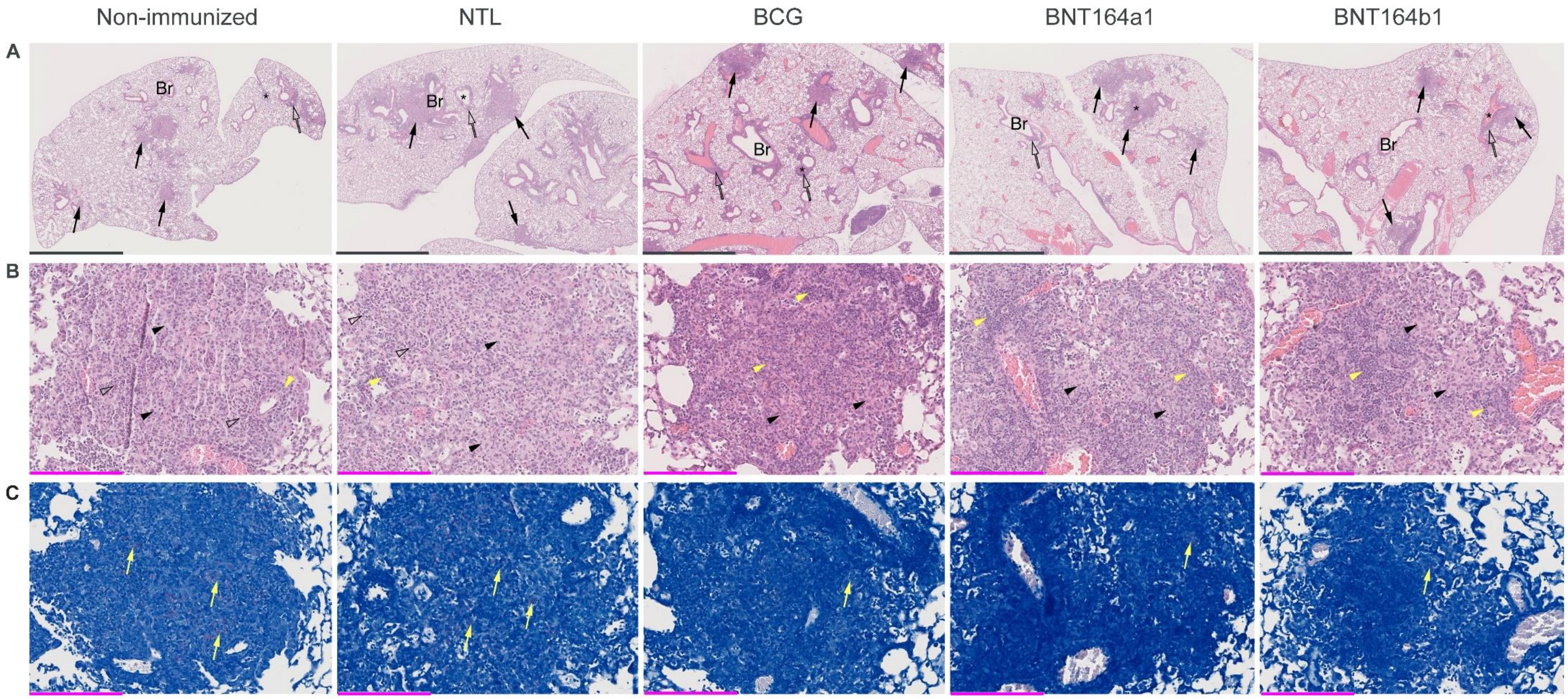


**Fig. S6. Pathology of *Mtb* HN878 infected mouse lungs**

The study design is depicted in Fig. 6A. Right lung lobes from four mice per group were paraffin embedded, formalin fixed and stained with hematoxylin & eosin (A-B) or acid-fast staining (C). Representative images are shown. (A) Closed black arrows indicate granulomas, while open black arrows indicate perivascular/peribronchiolar lymphocyte infiltrate cuffs surrounding either blood vessels (*) or bronchioles (Br). (B) Closed black arrowheads point to macrophages, yellow arrowheads point to lymphocytes, and open black arrowheads point to neutrophils. (C) Yellow arrows indicate acid-fast bacteria within granulomas. Black scale bars depict 2 mm (A), while pink scale bars depict 200 µm (B-C).

### Supplementary figure 7

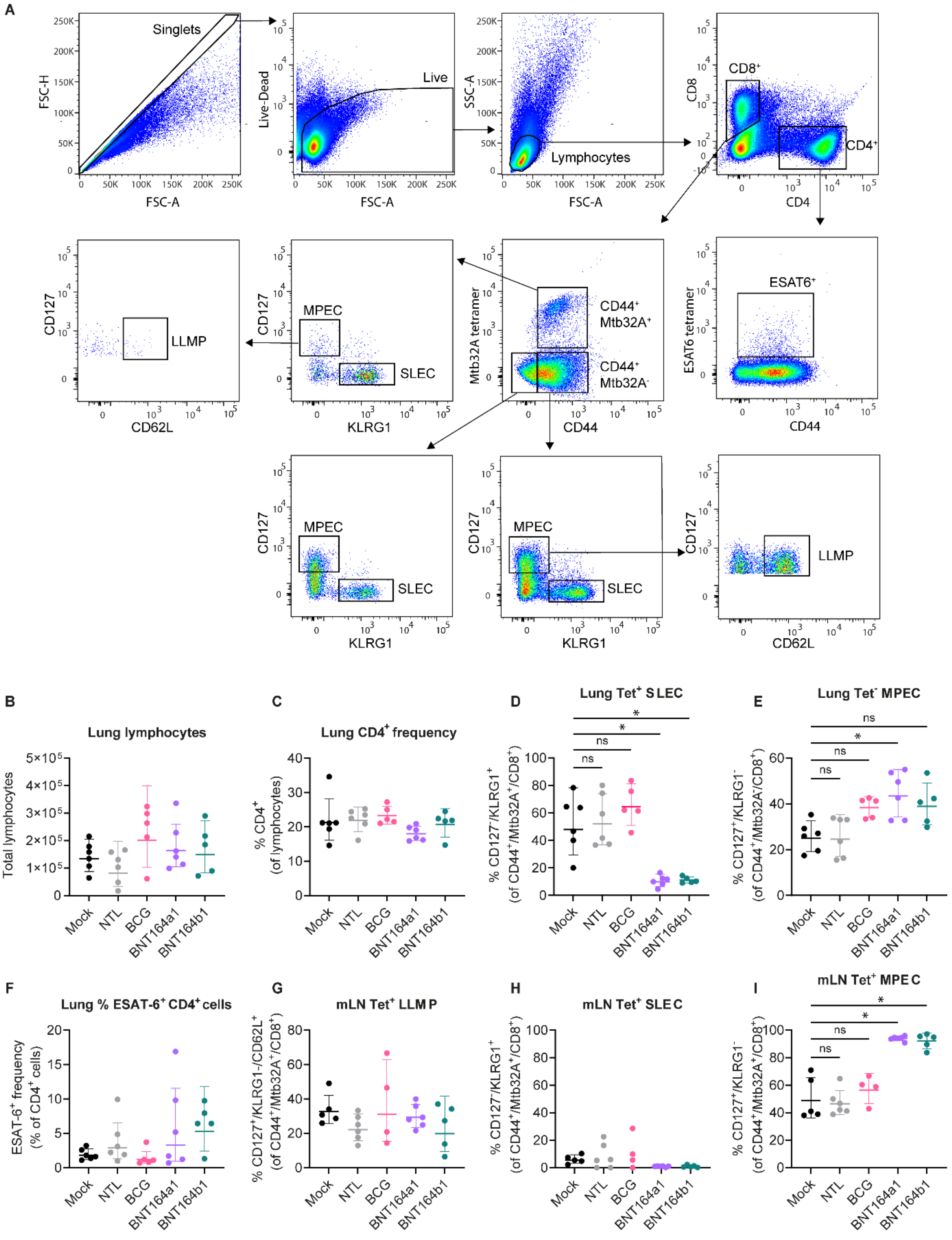


**Fig. S7. Lymphocyte quantification and phenotype in lungs and mesenteric lymph nodes of *Mtb* HN878-infected mice.**

The study design is depicted in Fig 6A. Right lung lobes (B–F), and mLNs (G–I) were homogenized 30 days post-infection and analyzed by flow cytometry. (A) Gating strategy shown on a lung sample of a saline-injected animal. All data was analyzed by Kruskal-Wallis non-parametric ANOVA followed by Dunn´s test for multiple comparisons against the saline control only if $p < 0.05$. Abbreviations: LLMP = Long-lived memory precursor; mLN = mesenteric lymph node; MPEC = Memory precursor effector cell; SLEC = Short-lived effector cell; ns = $p > 0.05$; *$p < 0.05$.
